# Studies on CRMP2 SUMOylation-deficient transgenic mice identify sex-specific NaV1.7 regulation in the pathogenesis of chronic neuropathic pain

**DOI:** 10.1101/2020.04.20.049106

**Authors:** Aubin Moutal, Song Cai, Jie Yu, Harrison J. Stratton, Aude Chefdeville, Kimberly Gomez, Dongzhi Ran, Cynthia L. Madura, Lisa Boinon, Maira Soto, Yuan Zhou, Zhiming Shan, Lindsey A. Chew, Kathleen E. Rodgers, Rajesh Khanna

**Affiliations:** Department of Pharmacology, College of Medicine, The University of Arizona, Tucson, Arizona, 85724 United States of America; Center for Innovation in Brain Sciences, University of Arizona, Tucson, Arizona 85721, United States of America; BIO5 Institute, University of Arizona, 1657 East Helen Street Tucson, AZ 85719; Department of Anesthesiology, College of Medicine, The University of Arizona, Tucson, Arizona, 85724 United States of America

## Abstract

The sodium channel NaV1.7 is a master regulator of nociceptive neuronal firing. Mutations in this channel can result in painful conditions as well as produce insensitivity to pain. Despite being recognized as a “poster child” for nociceptive signaling and human pain, targeting NaV1.7 has not yet produced a clinical drug. Recent work has illuminated the NaV1.7 interactome, offering insights into the regulation of these channels and identifying potentially new druggable targets. Amongst the regulators of NaV1.7 is the cytosolic collapsin response mediator protein 2 (CRMP2). CRMP2, modified at Lysine 374 (K374) by addition of a small ubiquitin-like modifier (SUMO), bound NaV1.7 to regulate its membrane localization and function. Corollary to this, preventing CRMP2 SUMOylation was sufficient to reverse mechanical allodynia in rats with neuropathic pain. Notably, loss of CRMP2 SUMOylation did not compromise other innate functions of CRMP2. To further elucidate the *in vivo* role of CRMP2 SUMOylation in pain, we generated CRMP2 K374A knock-in (CRMP2^K374A/K374A^) mice in which Lys374 was replaced with Ala. CRMP2^K374A/K374A^ mice had reduced NaV1.7 membrane localization and function in female, but not male, sensory neurons. Behavioral appraisal of CRMP2^K374A/K374A^ mice demonstrated no changes in depressive or repetitive, compulsive-like behaviors, and a decrease in noxious thermal sensitivity. No changes were observed in CRMP2^K374A/K374A^ mice to inflammatory, acute, or visceral pain. In contrast, in a neuropathic model, CRMP2^K374A/K374A^ mice failed to develop persistent mechanical allodynia. Our study suggests that CRMP2 SUMOylation-dependent control of peripheral NaV1.7 is a hallmark of chronic, but not physiological, neuropathic pain.

## 1. Introduction

The voltage-gated sodium channel NaV1.7 is a “poster child” target in pain signaling: gain-of-function mutations in the human NaV1.7 gene *SCN9A* can produce sensory neuron hyperexcitability associated with severe pain as well as insensitivity to pain [16]. While other sodium channels regulate the propagation of action potentials along nerves, NaV1.7 is upstream and defines the threshold at which an action potential will be elicited [44].

Alterations of NaV1.7 trafficking are important in the etiology of neuropathic pain. Mapping the NaV1.7 interactome has shed light on novel proteins involved in regulation of trafficking and degradation of NaV1.7 [10; 35]. In neuropathic pain, the expression of proteins regulating trafficking of voltage gated sodium channels (VGSCs) is dysregulated [4; 38]. In particular, upregulation of the VGSC β-subunits [4] and downregulation of Nedd4-2 (an E3 ubiquitin ligase)[38] following a spared nerve injury (SNI)[15], converge to functionally upregulate NaV1.7.

We identified the collapsin response mediator protein 2 (CRMP2) as a regulator of NaV1.7 function [17-19; 22; 48]. Our laboratory uncovered the logical coding of CRMP2’s cellular actions [11; 51]. The argument path underlying NaV1.7 regulation is defined by “IF CRMP2 is phosphorylated by cyclin-dependent kinase 5 (Cdk5) (S522) AND SUMOylated (K374) by E2-conjugating enzyme (Ubc9) THEN NaV1.7 is functional” [17; 51], “IF NOT SUMOylated, THEN CRMP2 recruits the endocytic proteins Numb, Nedd4-2, and Eps15” which will trigger clathrin mediated endocytosis of NaV1.7 [17]. This CRMP2 code posits that SUMOylation of CRMP2 is necessary for NaV1.7 membrane localization/function. In neuropathic pain, we reported a direct correlation between increased NaV1.7 function and CRMP2 SUMOylation [47]. Genetic interference with CRMP2 SUMOylation, via expression of a SUMO-null CRMP2 mutant (K374A), reversed mechanical allodynia in a rat model of neuropathic pain [47]. A “decoy” peptide that competitively interfered with CRMP2’s interaction with Ubc9 also resulted in decreased CRMP2 SUMOylation and NaV1.7 currents which reversed mechanical allodynia in rats with neuropathic pain [22; 48]. Importantly, deletion of CRMP2 SUMOylation did not impair its other functions (e.g., CaV2.2 currents or neurite outgrowth [19; 51; 73]). These studies revealed CRMP2 SUMOylation as a specific regulatory mechanism underlying NaV1.7’s pathological function in neuropathic pain. Overall, our work has established that interfering with CRMP2 SUMOylation is a safe and efficient therapeutic approach for disrupting neuropathic pain [11; 22; 47].

To further elucidate the *in vivo* role of the SUMOylation of CRMP2 at Lys374 in pain, we generated CRMP2 K374A knock-in (CRMP2^K374A/K374A^) mice in which Lys374 was replaced with Ala. The CRMP2^K374A/K374A^ mice had reduced NaV1.7 membrane localization and function in female, but not male, sensory neurons. Behavioral appraisal of the CRMP2^K374A/K374A^ mice demonstrated no changes in depressive or repetitive, compulsive-like behaviors, and a decrease in thermal nociception. No changes were observed in CRMP2^K374A/K374A^ mice subjected to inflammatory (formalin), acute (post-surgical), or visceral (acetic-acid evoked writhing) pain models. In contrast, in the partial denervation of SNI, CRMP2^K374A/K374A^ mice failed to develop persistent mechanical allodynia. Our study suggests that CRMP2 SUMOylation-dependent control of NaV1.7 engages a novel nociceptive pathway related to the transition to chronic neuropathic pain.

## 2. Materials and Methods

### 2.1. Ethics Statement

All animal use was conducted in accordance with the National Institutes of Health guidelines, and the study was carried out in strict accordance with recommendations in the Guide for the Care and Use of Laboratory Animals of the University of Arizona (Protocol #: 16-141). All animals were housed and bred in the University of Arizona Laboratory Animal Research Center. Mice were housed in groups of 4-5 in a dedicated housing facility with ad libitum access to food and water on a 12-hour light/dark cycle. The same mice were tested using no more than two non-invasive behavioral assessments, reducing the total number of mice needed to complete the study. All efforts were made to minimize animal suffering to reduce the number of the animals used. All behavioral experiments were performed by experimenters who were blinded to the genotype, and sex, and treatment groups.

### 2.2. Reagents

All chemicals, unless noted were purchased from Sigma (St. Louis, MO). PF-05089771 [2,2-diphenyl-*N*-(4-(*N*-thiazol-2-ylsulfamoyl)phenyl)acetamide] was purchased from Biotechne - R&D Systems (Minneapolis, MN). Pitstop2 (Cat# ab120687) was from Abcam (Cambridge, MA). Validated antibodies were purchased as follows: anti-CRMP2 polyclonal antibody (Sigma-Aldrich Cat# C2993 Research Resource Identifier (RRID): AB_1078573), anti-NaV1.7 (Cat# 75-103, NeuroMab, Davis, CA, RRID: AB_2184355), SUMO1 (Sigma-Aldrich Cat# S8070 RRID:AB_477543) [17; 19].

### 2.3. Generation, genotyping, and general health of CRMP2^K374A/K374A^ mice

CRISPR guide RNAs (gRNAs), used for mouse *dpysl2* gene (Gene ID: 12934) K374A knock-in were designed using CRISPR.mit.edu [12]. Two guides RNAs were selected and produced by PCR using the pX330 plasmid (Addgene, Watertown, MA) as a template. The forward primer consisted of T7 promoter sequence and the target sequence. sgRNAs (single guide RNAs) were designed to target the following sequences in the *dpysl2* gene: GTGTACATGCTCTGTCCTGCAGG (gRNA1), TGCTCTGTCCTGCAGGTCACTGG (gRNA2) (**Figure 1A**). gRNAs were made by in vitro transcription using Ambion’s MEGAshortscript kit and afterwards purified with MEGAclear kit. Commercial Cas9 mRNA was ordered from TriLink (L-6125, San Diego, CA) and used for the embryo microinjection. Cas9 protein was ordered from PNA Bio (CP01-50, Thousand Oaks, CA) and used to check cutting efficiency of the guides in vitro. ssODNs with 60-70 bp homology to sequences on each side of each gRNA-mediated double-stranded break were designed and ordered from IDT (Coralville, IA). A silent mutation introducing *MspI* restriction site in the oligo was created for genotyping purposes.

**Figure 1.**
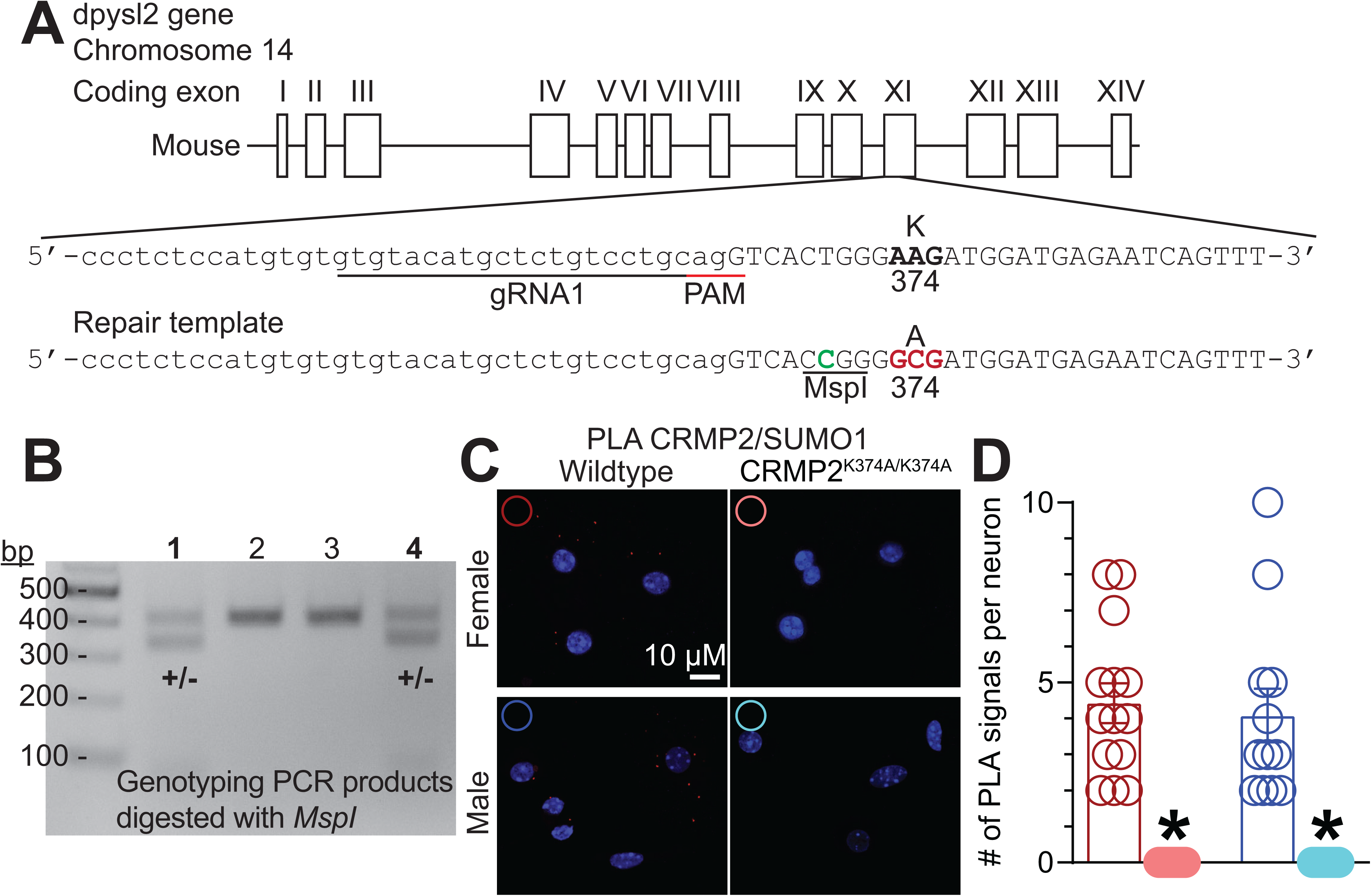
Generation of CRMP2^K374A/K374A^ knock-in mice. (**A**) Strategy for the generation of the CRMP2– K374A knock-in mutation in mouse ES cells. Guide RNA (gRNA) design for targeting the mouse *dpysl2* locus. The protospacer-adjacent motif (PAM) sequence is in underlined in red. The AAG to GCG mutation is in bold font. (**B**) PCR results of founder mice. Mice from two litters were genotyped by PCR amplification and MspI digestion. MspI digestion products were electrophoresed on a 1.5% agarose gel. Wild type mouse genomic DNA was used as negative control (not shown). (**C**) Representative images of mouse DRG cultures following proximity ligation assay (PLA) between CRMP2 and SUMO1. The PLA immunofluorescence labeled sites of interaction between CRMP2 and SUMO1 (red puncta). Additionally, nuclei are labeled with the nuclear labeling dye 4’,6-diamidino-2-phenylindole (DAPI). Scale bar: 10 µm. (**D**) Quantification of PLA puncta per neuron show that in DRGs from CRMP2^K374A/K374A^ mice, the number of CRMP2-SUMO1 interactions are significantly reduced compared to wildtype DRGs (Kruskal-Wallis test with Dunn’s multiple comparison post-hoc test: p<0.0001 comparing female WT vs. female CRMP2^K374A/K374A^; p<0.0001 comparing male WT vs. male CRMP2^K374A/K374A^; and p>0.9999 comparing male WT vs. female WT). Error bars indicate mean ± SEM from 12-16 cells.

Fertilized eggs were collected from the oviducts of super ovulated C57BL6/J females. Microinjection was performed by continuous flow injection of the Cas9/gRNA/ssODN mixture into the pronucleus of 1-cell zygotes using the following final concentrations 100/50/200 ng/µl respectively.

Tail-tipping of the newborn mice was utilized to purify DNA for genotyping by PCR, employing two screening primers: forward, 5’-AGGGAAATCCCAGGATGAAT and reverse, 5’-CGGGGGTAAAGGTTGAAGAC, producing 403bp band for the wild type and two additional bands of 329bp and 74bp in the positive mice when restricted with *MspI*. gRNA1 and the corresponding oligonucleotides were successful in introducing the desired mutation.

The CRMP2^K374A/K374A^ mice appeared healthy and showed no obvious differences in physical characteristics. During breeding and until weaning of the pups, mice were fed three times weekly with Love Mash™ Rodent Reproductive diet (Cat# S3823P, Bio-Serv, Flemington, NJ) to improve fertility for both males and females, increase litter size and increase pup survival rate. There were no significant differences in body weight within sexes of CRMP2^K374A/K374A^ mice compared to their WT littermates (6–22 weeks of age) – Females: 21.0 ± 0.61 g in WT [n = 24]; 20.0 ± 0.29 g in CRMP2^K374A/K374A^ [n = 32]; Males: 28.0 ± 0.71 g in WT [n = 34]; 27.0 ± 0.49 g in CRMP2^K374A/K374A^ [n = 42] (p>0.9999, Kruskal-Wallis with Dunn’s post hoc test) (**Supplementary Figure 1**). Male mice, irrespective of genotype, were larger than female mice (p<0.0001, Kruskal-Wallis with Dunn’s post hoc test) (**Supplementary Figure 1**). Analyses of heterozygous CRMP2 K374A knock-in mice in all assays revealed them to be no different than wildtype mice (data not shown); for clarity, only WT and homozygous CRMP2^K374A/K374A^ mice are shown.

### 2.4. Preparation of acutely dissociated dorsal root ganglia neurons from wildtype (WT) and CRMP2^K374A/K374A^ mice

Wildtype and CRMP2^K374A/K374A^ mice were deeply anaesthetized with isoflurane overdose (5% in oxygen) and sacrificed by decapitation. Dorsal root ganglia (DRG) were quickly removed, trimmed at their roots, and enzymatically digested in 3 mL bicarbonate-free, serum-free, sterile DMEM (Cat# 11965, Thermo Fisher Scientific, Waltham, MA) solution containing neutral protease (1.87 mg/ml, Cat#LS02104; Worthington, Lakewood, NJ) and collagenase type I (3 mg/mL, Cat# LS004194, Worthington, Lakewood, NJ) and incubated for 60 minutes at 37°C under gentle agitation. Dissociated DRG neurons (∼1.5 × 10^6^) were then gently centrifuged to collect cells and washed with DRG media (DMEM containing 1% penicillin/streptomycin sulfate from 10,000 μg/mL stock, and 10% fetal bovine serum (Hyclone)) before plating onto poly-D-lysine– and laminin-coated glass 12-mm glass coverslips. The experiments were performed within 48 h of plating DRG neurons since electrophysiological profiles did not undergo changes within this period.

### 2.5. Immunofluorescence, confocal microscopy, and quantification of NaV1.7

Immunofluorescence was performed on wildtype and CRMP2^K374A/K374A^ mice DRGs as described previously [17]. Briefly, cells were fixed using ice cold methanol for 5 minutes and allowed to dry at room temperature. Cells were rehydrated in PBS and anti-NaV1.7 antibody (1/1000) was added in PBS with 3% BSA for 1 hour at room temperature. Cells were then washed three times in PBS and incubated with PBS containing 3% BSA and secondary antibodies (Alexa 488 goat anti-mouse; Thermofisher) for 1 hour at room temperature. After washing with PBS, cells were stained with 4’,6-diamidino-2-phenylindole (DAPI, 50µg/ml) and mounted in Fluoro-gel (Cat# 17985, electron microscopy sciences). Immunofluorescent micrographs were acquired using a plan-Apochromat 6320x/0.8 objective on a Zeiss LSM880 confocal microscope operated by the Zen Black software (Zeiss). Camera gain and other relevant settings were kept constant. Membrane immunoreactivity was calculated by measuring the signal intensity in the area contiguous to the boundary of the cell. Membrane to cytosol ratio was determined by defining regions of interest in the cytosol and on the membrane of each cell using Image J. Total fluorescence was normalized to the area analyzed and before calculating the ratios.

### 2.6. Proximity ligation assay (PLA)

PLA was performed as described previously [48-50; 52] to visualize protein-protein interactions using microscopy. This technique is based on paired complementary oligonucleotide-labeled secondary antibodies that can hybridize and amplify a red fluorescent signal only when bound to two corresponding primary antibodies whose targets are in close proximity (i.e., within ∼30 nanometers). DRG neurons were fixed using ice cold methanol for 5 minutes and allowed to dry at room temperature. Primary antibodies (1/1000) were incubated for 1 hour at RT in PBS with 3% BSA before 3 washes in PBS for 5 minutes at room temperature. The proximity ligation reaction and amplification of signal were performed according to the manufacturer’s protocol using the Duolink Detection Kit with PLA PLUS and MINUS probes for mouse and rabbit antibodies (Duolink, Sigma). DAPI stain was used to detect cell nuclei. Immunofluorescent micrographs were acquired using a plan-Apochromat 6320x/0.8 objective on a Zeiss LSM880 confocal microscope operated by the Zen Black software (Zeiss). Image J was used to count the number of PLA puncta per cell.

### 2.7. Whole-cell electrophysiological recordings of sodium, calcium and potassium currents in acutely dissociated DRG neurons from WT and CRMP2^K374A/K374A^ mice

All recordings were obtained from acutely dissociated DRG neurons from genotyped WT and CRMP2^K374A/K374A^ mice, using procedures adapted from those as described in our previously published work [19; 32; 48]; the electrophysiologists were blinded to the genotype and sex.

Sodium currents: the internal solution consisted of (in mM): 140 CsF, 10 NaCl, 1.1Cs-EGTA, and 15 HEPES (pH 7.3, mOsm/L = 290-310) and external solution contained (in mM): 140 NaCl, 30 tetraethylammonium chloride, 10 D-glucose, 3 KCl, 1 CaCl_2_, 0.5 CdCl_2_, 1 MgCl_2_, and 10 HEPES (pH 7.3, mOsm/L = 310-315). DRG neurons were interrogated with current-voltage (I-V) and activation/inactivation voltage protocols as previously described [17; 24]. The voltage protocols were as follows: (a) I-V protocol: from a −60 mV holding potential, cells were depolarized in 150-millisecond voltage steps from −70 to +60 mV (5-mV increments) which permitted acquisition of current density values such that we could analyze activation of sodium channels as a function of current vs voltage and infer peak current density (normalized to cell capacitance (in picofarads, pF)), which occurred between ∼0 to 10 mV; (b) inactivation protocol: from a −60 mV holding potential, cells were subjected to hyperpolarizing/repolarizing pulses for 1 second between −120 to 0 mV (+10 mV steps). This increment conditioned various proportions of channels into a state of fast-inactivation – in this case 0-mV test pulse for 200 milliseconds was able to reveal fast inactivation when normalized to maximum sodium current. Because of the differential inactivation kinetics of TTX-resistant and TTX-sensitive channels, the fast inactivation protocol allowed subtraction of electrically isolated TTX-R (current available after –40mV prepulse) from total current (current available after −120mV prepulse), as previously described [17]. Pipettes with 1 to 3 MΩ resistance were used for all recordings. In experiments where clathrin mediated endocytosis was prevented with 20 µM Pitstop2 [70], the compound was added on the cells 30 minutes prior to the experiment.

To isolate calcium currents (I_Ca_), Na^+^ and K^+^ currents were blocked with 500 nM TTX (Alomone Labs) and 30 mM tetraethylammonium chloride (Sigma). Extracellular recording solution (at ∼310 mOsm) consisted of the following (in mM): 128 N-methyl-*D*-glucamine, 10 BaCl_2_, 5 KCl, 2 MgCl_2_ 10 tetraethylammonium chloride, 10 Na-HEPES, 10 D-glucose, pH at 7.4; with 1 μM TTX, and 10 μM nifedipine. The intracellular recording solution (at ∼310 mOsm) consisted of the following (in millimolar): 150 CsCl_2_, 10 HEPES, 5 Mg-ATP, 5 BAPTA, at pH 7.4. Activation of I_Ca_ was measured using a holding voltage of −90 mV with 200-ms voltage steps applied at 5-s intervals in +10 mV increments from −70 to +60 mV. Current density was calculated as peak I_Ca_ divided by cellular capacitance. Steady-state inactivation of I_Ca_ was determined by applying an 1500 ms conditioning prepulse (−100 to +30 mV in +10 mV increments) after which, the voltage was stepped to +10 mV for 200-ms; a 15-s interval separated each acquisition.

To isolate potassium currents (I_k_), DRG neurons were bathed in buffer composed of (in mM): 140 N-methyl-glucamine chloride, 5 KCL, 2 CaCl_2_, 1 MgCl_2_, 10 HEPES, and 10 glucose, pH adjusted to 7.4 with KOH. Recording pipettes were filled with internal solution (in mM): 140 KCL, 5 MgCl_2_, 4 ATP, 0.3 GTP, 2.5 CaCl_2_, 5 EGTA, and 10 HEPES, adjusted to pH 7.3 with KOH. Neurons were then subjected to activation I_k_, fast inactivating I_k_A, slow inactivating IkS and inactivation I_k_A/I_k_S protocols as previously described [48]. Membrane holding was set to –60 mV. I_k_ activation was determined by applying 300 ms incremental voltage steps at 5 second intervals from −80 to +60 mV (in + 20 mV increments), after a 4-s prepulse to −100mV. I_k_S was recorded in response to incremental voltage steps of 500 milliseconds, applied at 5 second intervals from - 80 to +60 mV (in + 20-mV increments), after a 4-s prepulse to −40mV.The fast inactivating I_k_A was inferred by digitally subtracting the currents from the prepulse to –100mV to the prepulse to –40mV. Inactivation of I_k_A was determined using a series of 4-second prepulses that ranged from –100 to –40 mV with an increments of +10-mV per step that were immediately followed by a 200-millisecond step to +60 mV.

### 2.8. Preparation of spinal cord slices for slice-electrophysiological recording

As described previously [75], age-matched genotyped WT and CRMP2^K374A/K374A^ mice (postnatal 12-15 days) were deeply anesthetized with isoflurane (4% for induction and 2% for maintaining). For spinal nerve blocking, 0.1-0.2 mL of 2% lidocaine was injected to both sides of L4 to 5 lumbar vertebrae. Laminectomy was performed from mid-thoracic to low lumbar levels, and the spinal cord was quickly removed to cold modified ACSF oxygenated with 95% O_2_ and 5% CO_2_. The ACSF for dissection contained (in millimolar): 80 NaCl, 2.5 KCl, 1.25 NaH_2_PO_4_, 0.5 CaCl_2_.2H_2_O, 3.5 MgCl_2_.6H_2_O, 25 NaHCO_3_, 75 sucrose, 1.3 ascorbate, 3.0 sodium pyruvate, with pH at 7.4 and osmolarity at 310 mOsm. Transverse 360 – µm thick slices were obtained by a vibratome (VT1200S; Leica, Nussloch, Germany). Slices were then incubated for 40 minutes at 37°C before resting for at least 1 hour at RT in an oxygenated recording solution containing (in millimolar): 125 NaCl, 2.5 KCl, 2 CaCl_2_.2H_2_O, 1 MgCl_2_.6H_2_O, 1.25 NaH_2_PO_4_, 26 NaHCO_3_, 25 D-glucose, 1.3 ascorbate, 3.0 sodium pyruvate, with pH at 7.4 and osmolarity at 320 mOsm. The slices were then positioned in a recording chamber and continuously perfused with oxygenated recording solution at a rate of 3 to 4 mL/min before electrophysiological recordings at room temperature.

#### 2.8.1. Whole-cell patch recordings of spontaneous excitatory post-synaptic currents in substantia gelatinosa neurons of spinal cord slices

Substantia gelatinosa neurons were visualized and identified in the slices by means of infrared DIC video microscopy on a FN1 Nikon upright microscope (Nikon, Tokyo, Japan) outfitted with a 3 40/0.80 objective (water-immersion) and a charge-coupled device camera. Patch pipettes with resistance ranging from 6 to 10 MΩ were fabricated from borosilicate glass (Sutter Instruments, Novato, CA) on a 4-step micropipette P-90puller (Sutter Instruments, Novato, CA). The internal solution contained the following (in mM): 120 potassium gluconate, 20 HEPES, 20 KCl, 2 MgCl_2_.6H_2_O, 2 Na2 -ATP, 0.5 Na-GTP, 0.5 EGTA, with pH at 7.4 and osmolarity at 310 mOsm. The membrane was held at −60 mV using Patchmaster.

In voltage-clamp mode, whole-cell configurations were obtained. To record spontaneous excitatory postsynaptic currents (sEPSCs), bicuculline methiodide (10 μM) and strychnine (1 μM) were added to the recording solution to block γ-aminobutyric acid-activated (GABA) and glycine-activated (GlyR) currents, respectively. Hyperpolarizing step pulses (5 mV in intensity, 50 ms) were periodically delivered to test the access resistance (15-25 MΩ), and recordings were stopped if the access resistance changed >20%. For each recording, sEPSCs were acquired for a duration of 2 minutes. Currents were filtered and digitized at 3 kHz and 5 kHz, respectively. Data were evaluated by the Mini-Analysis (Synatosoft Inc, NJ) and Clampfit 10.7 Program. The amplitude and frequency of sEPSCs were compared between neurons from different sex and genotype groups. All data were collected in 1 hour within drug perfusion.

### 2.9. Assessment of long-term potentiation (LTP) recordings in hippocampi of WT and CRMP2^K374A/K374A^ mice

Male and female mice between 6-9 weeks of age were used for this experiment. For preparation of coronal brain slices containing the hippocampus, mice were first anesthetized using isoflurane then swiftly decapitated. The brain was extracted, taking care to remove the meninges, and immediately placed in ice-cold sucrose based cutting buffer with the following composition (in mM) 212.7 Sucrose, 5 KCl, 1.25 NaH_2_PO4, 26 NaHCO_3_, 3 MgSO_4_, 1 CaCl_2_, 10 Glucose. The frontal portion of the cortex and the cerebellum were then removed using a razor blade to allow the brain to be fixed to the cutting stage of a Leica VT1200S vibratome (Leica, Nussloch, Germany) with cyanoacrylate adhesive. The stage was fully immersed in sucrose based artificial cerebrospinal fluid (aCSF) that was continuously bubbled with carbogen (95% O_2_/5% CO_2_) and the blade was lowered to a cutting angle of 18 degrees relative to the horizontal axis of the vibratome. The blade was advanced at 0.18 mm/s with a swing amplitude of 1.3 mm and the brain was cut into 400 µm thick sections. Brain slices were subsequently cut in half along the sagittal midline and transferred to a holding chamber (Brain Slice Keeper, AutoMate Scientific) at room temperature (25°C) that contained regular aCSF and was continuously bubbled with carbogen. The recording aCSF contained the following (in mM): 119 NaCl, 2.5 KCl, 1.0 NaH_2_PO_4_, 26.2 NaHCO_3_, 1.3 MgSO_4_, 2.5 CaCl_2_, 11 Glucose. The slices were left to recover for at least 1 hour in this chamber before being transferred to a multi-electrode array (MEA P515A, MED 64 AlphaMed Scientific, Japan) for recording of hippocampal field excitatory post synaptic potentials.

The recording electrodes in the MEA system are composed of indium-tin oxide in a polyimide substrate that were then coated with carbon nanotubes to reduce electrical impedance and increase surface area of the electrodes. Once in the recording chamber the slice was fixed with a thin mesh and a U-shaped harp anchor (Warner Instruments) to prevent motion during recording. Fresh aCSF was bubbled with carbogen and warmed to 30°C then delivered to the recording chamber using a peristaltic pump at a rate of approximately 2 mL/min. The solution level in the recording chamber was maintained at the interface condition by adjusting the height of the inflow and outflow tubes. Humidified carbogen was delivered to the recording chamber to provide an oxygenated environment. The slice was positioned over the recording array using an inverted microscope such that the pyramidal cell layer of region CA1 of the hippocampus was parallel to one of the rows of recording electrodes. An electrode at the edge of the array located along the Schaffer Collateral projections from region CA3 to region CA1 was then selected as the stimulation electrode based on the response of the slice to a test pulse of 15 mA. Once a suitable electrode was identified, this single electrode was used for all subsequent stimulation during the recording period. The slice was left to equilibrate to the recording chamber conditions for approximately 20 minutes before recording commenced.

Next, the input output relationship was determined for each slice beginning with a stimulation intensity of 10 mA and continuing in 5 mA steps delivered at 0.033 Hz until a maximal response was achieved. For all remaining protocols, the stimulation intensity was set to 50% of this maximal value. Paired pulse responses were then collected with pulse intervals of 15, 50, 150 and 400 ms and at least 2 minutes between each recording to prevent unintended potentiation. Subsequently, a baseline period of 20 minutes was recorded with stimulation occurring at 0.033 Hz. Any slice where the baseline fEPSP slope value varied by more than ± 10% was rejected as unstable and not included for analysis. After the 20-minute baseline period, theta-burst stimulation (TBS), consisting of 5 bi-phasic current pulses at 100 Hz repeated 10 times in 2 seconds, was used to elicit long term potentiation (LTP) in region CA1. The slope and amplitude of fEPSP responses were recorded for an additional 60 minutes following LTP induction with stimulation pulses delivered at 0.033 Hz. Responses that began to drift following stimulation were excluded from data analysis. To determine whether the input output functions were different between groups, linear regression was used to generate a linear best fit function for both sets of data and an analysis of covariance was used to determine whether the slope of each function was significantly different [29]. The paired pulse ratio was analyzed using a two-way ANOVA with genotype and time as factors. The level of potentiation was calculated using an average of the fEPSP slope over the last five minutes of the recording period and a two-tailed t-test to determine whether the level of potentiation was significantly different between groups.

### 2.10. Assessment of potential CNS side effects

#### 2.10.1 Marble burying test as a measure of compulsive-like behaviors

The marble-burying test [6]was used to evaluate compulsive-like behaviors in WT and CRMP2^K374A/K374A^ mice. Each cage was filled with 5-cm of bedding chips lightly pressed on the floor to create a flat, even surface. The dimensions of the testing cage were 15-cm ×30-cm ×15-cm and were identical to the housing cages. Mice were habituated to the experimental room for 30 minutes before the beginning of the experiment. A total of 17 glass marbles (1.5 cm in diameter) were evenly distributed on the bedding surface. Mice were placed in the cages (1 mouse/cage) and allowed to behave freely for 30 minutes. After 30 minutes, the mice were returned to their home cages and the number of marbles at least 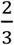 buried orcovered with bedding were counted.

#### 2.10.2. Nestlet shredding test as a measure of compulsive-like behaviors

The nestlet shredding test [57] was also used to evaluate compulsive-like behaviors in WT and CRMP2^K374A/K374A^ mice. The testing cages, bedding and nestlet used in this test are the same as those used in home cages for the mice. The dimensions of the testing cage were 15-cm x30-cm x15-cm and were identical to the housing cages. Mice were habituated to the experimental room for 30 minutes before the beginning of the experiment. The floor of each cage was lightly covered with bedding. A nestlet was weighed, then placed in the center of the cage. Mice were placed in the cages (1 mouse/cage) and allowed to behave freely for 30 minutes. After 30 minutes, the animals were returned to their home cages. The nestlet was dried then weighed, yielding the change in nestlet weight. The percentage of weight reduction was used as an indication of compulsory-shredding.

#### 2.10.3. Tail suspension test (TST) as a measure of depression-associated behavior

The TST was conducted according to standard procedures [13]. Briefly, each mouse was suspended by the tail at a height of 40 cm by taping the rail to a horizontal bar so that the tail was perpendicular to the bar. Behavior was video recorded for 5-min and later analyzed by an experimenter who was blinded to the groups. An animal was considered to be immobile when it did not show any movement and hung passively. If a mouse climbed its tail the mouse was gently pulled back down and the trial continued. Mice that climbed their tails during more than 20% of the trial (i.e. >60 seconds) were eliminated from the final analysis. Duration and frequency of immobility were the main parameters measured.

#### 2.10.4. Elevated plus maze (EPM) test as a measure of anxiety-associated behavior

The EPM consists of four elevated (50 cm) arms (50 cm long and 10 cm wide) with two opposing arms constructed of 30 cm high opaque walls. EPM testing occurred in a quiet testing room with ambient lighting at ∼500 lux. On the day of testing, mice were acclimated to the testing room for 20 minutes. Each mouse was placed in a closed arm, facing the platform entrance and all cage mates were started in the same closed arm. Each mouse was allowed 5 minutes to explore the EPM and then returned to its home cage. Between animals, the EPM was cleaned thoroughly with Versa-Clean (Fisher Scientific). EPM performance was recorded using an overhead video camera (MHD Sport 2.0 WiFi Action Camera, Walmart.com) for quantification and scoring. Entry into the open and closed arm were defined as the front two paws entering the arm. The open arm time began the moment the front paws entered the open arm and ended upon exit.

### 2.11. Hot plate test

Wildtype and CRMP2^K374A/K374A^ mice were placed on a metal plate (Stoelting, Wood Dale, IL) warmed to 48°C, 52°C or 55°C and a timer was started. Latency to the first response (flinching or licking the hind paws or jumping) was recorded. Cut-off times were used to prevent tissue damage (60 s, 20 s and 15 s, respectively) and any mouse that reached the cut-off time was assigned the value of the cut-off time as their latency time.

### 2.12. Tail-flick test

The distal third of the tail of wildtype and CRMP2^K374A/K374A^ mice was immersed in warm water (52°C). Latency to remove the tail from the water (tail flick latency) was recorded. To prevent tissue damage, a 10-s cut-off was used and animals that did not react before the cut-off were attributed a latency of 10 s.

### 2.13. Formalin test

A 2% formalin solution in saline was injected into the plantar skin of left hindpaw of the mouse and spontaneous nociceptive responses were recorded and analyzed. The duration of licking behaviors was counted for 1-h in 5-min intervals. The first phase of nociceptive responses was analyzed during 0–10 min, and the second phase of nociceptive responses was analyzed during 10–60 min.

### 2.14. Acetic acid-induced writhing test

The acetic acid-induced writhing test is a model of visceral pain where a chemical irritant -acetic acid- is injected intraperitoneally, inducing abdominal contractions and extension of back limbs, collectively defined as ‘*writhes*’ [41]. Mice were first allowed to acclimate for 15-min within the clear Plexiglas chamber used for the experiment or until exploratory behavior ceased. Mice were then immediately injected in the peritoneum of the left lower quadrant of the abdomen with 300 μL of 1% acetic acid in normal saline using a 27-gauge needle. Mice were placed in the testing chamber after the acetic acid injection and their subsequent nociceptive behavior was recorded and scored (in an investigator blinded manner): writhing behavior was assessed between 5 and 25 min after the injection.

### 2.15. Paw incision model of postoperative pain

A mouse model of surgical pain was generated by plantar incision as previously described [59]. Age-matched and genotyped WT and CRMP2^K374A/K374A^ mice were anesthetized with isoflurane vaporized through a nose cone. The plantar aspect of the left hind paw was scrubbed with betadine and 70% alcohol three times. A 1-cm long incision, starting 0.5 cm from the heel and extending toward the toes, was made with a number 11 blade, through the skin and fascia of the plantar aspect of the left hind paw including the underlying muscle. The plantaris muscle was then elevated and abraded to produce injury but leaving the muscle origin and insertion intact. After hemostasis with gentle pressure, the skin was closed with 2 mattress sutures of 6-0 nylon on a curved needle. Animals were allowed to recover for 24 hours and paw withdrawal thresholds were measured at 24 hours after surgery.

### 2.16 Veratridine model of acute pain

The lipophilic small molecule veratridine binds at the batrachotoxin/local anesthetic site common to all sodium channels, causing the channels to remain open upon repolarization following a step depolarization [8], and has been demonstrated to induce pain-like lifting and licking behaviors in mice [26]. Mice were placed in Plexiglas chambers (dimensions: 10 cm diameter, 14 cm tall cylinder with lid) for at least 30 minutes before testing commenced. Mice were briefly removed from their cylinder, restrained by hand, and 50 µl (1 mg) of veratridine dissolved in 0.1% DMSO in saline was administered as an intraplantar injection in the left hind paw. A video camera recorded activity for 20 minutes after the injection and digital video files were later scored in 5-min intervals for the total time spent licking the left hind paw.

### 2.17. Spared nerve injury (SNI) model of neuropathic pain

Age-matched and genotyped WT and CRMP2^K374A/K374A^ mice were habituated in plastic chambers on a mesh floor. Calibrated von Frey filaments with sequentially increasing spring coefficients were applied to the hind paw of each mouse, which allowed for the consistent application of constant force stimuli. One filament was applied 5 times in a round of testing. The filament force evoking paw withdrawal more than 3 times in a round of testing was defined as the mechanical threshold. The cutoff threshold was 4 g. To prepare the neuropathic pain model induced by spared nerve injury (SNI), we transected the common peroneal and tibial branches of the right sciatic nerve with ∼1 mm of nerve removed and left the sural nerve intact. The von Frey test was performed on the lateral part of the right plantar surface where the sural nerve innervates the hindpaw.

### 2.18. Flow cytometry analyses of brain, spinal cord, spleen, and blood of wildtype and CRMP2^K374A/K374A^ mice

#### 2.18.1 Brain and spinal cord cellular makeup

At necropsy the brain and spinal cord from genotyped WT and CRMP2^K374A/K374A^ mice were harvested and washed with PBS+2%FBS. Both tissues were then dissociated using the Adult Mouse Brain Dissociation kit (Miltenyi Biotech; Auburn, CA) as per the manufacturer’s specifications. After dissociation the single cell suspension was stained using antibodies to detect; CD45 (REA737, Miltenyi Biotech), CD11b (REA592, Miltenyi Biotech), CD31 (REA784, Miltenyi Biotech), GLAST (ACSA-1, Miltenyi Biotech), and O4 (REA576, Miltenyi Biotech). Stained samples were then analyzed on a MACSQuant 10 (Miltenyi Biotech) and data was processed on FlowLogic 7.3 (Inivai Technologies).

#### 2.18.2. Peripheral immunophenotyping

At necropsy the blood and spleen from genotyped WT and CRMP2^K374A/K374A^ mice were harvested and washed with PBS + 2%FBS. The spleen was dissociated using the Spleen Dissociation kit (Miltenyi Biotech; Auburn, CA) as per the manufacturer’s specifications. Both samples were then treated with RBC lysis buffer (Miltenyi Biotech; Auburn, CA) as per manufacturers specifications. The single cell suspension was stained using antibodies to detect; CD45(REA737, Miltenyi Biotech), CD3 (REA641, Miltenyi Biotech), CD4 (REA604, Miltenyi Biotech), CD8a (REA601, Miltenyi Biotech), CD69 (REA937, Miltenyi Biotech), and CD25 (REA568, Miltenyi Biotech). Stained samples were then analyzed on a MACSQuant 10 (Miltenyi Biotech) and data was processed on FlowLogic 7.3 (Inivai Technologies, Australia).

### 2.19. Statistical analyses

All data was first tested for a Gaussian distribution using a D’Agostino-Pearson test (Graphpad Prism 8 Software, Graphpad, San Diego, CA). The statistical significance of differences between means was determined using a parametric ANOVA followed by Tukey’s post-hoc test or a non-parametric Kruskal Wallis test followed by Dunn’s post-hoc test depending on if datasets achieved normality. Behavioral data with a time course were analyzed by Two-way ANOVA with Sidak’s post-hoc test. Differences were considered significant if p≤ 0.05. Error bars in the graphs represent mean ± SEM. See statistical analysis described in **Table 2**. All data were plotted in Prism 8.

**Table 1.** Gating properties of sodium currents recorded from DRG neurons of female and male wildtype and CRMP2^K374A/K374A^ knock-in mice^a^

|  | Females |  | Males |  |
| --- | --- | --- | --- | --- |
|  | Wildtype (WT) | CRMP2<sup>K374A/K374A</sup> | Wildtype (WT) | CRMP2<sup>K374A/K374A</sup> |
| **DMSO (Control)** |  |  |  |  |
| Activation |  |  |  |  |
| $V_{1/2}$ | -6.9±1.1(13) | -8.8±1.6(13) | -18.6±0.8(10) | -18.3±0.8(10) |
| $k$ | 7.6±1.0(13) | 9.2±1.5(13) | 5.2±0.8(10) | 5.8±0.7(10) |
| Inactivation |  |  |  |  |
| $V_{1/2}$ | -45.2±2.1(13) | -51.7±3.4(13) | -50.7±3.0(10) | -54.1±1.7(18) |
| $k$ | -12.9±2.1(13) | -16.4±3.8(13) | -17.0±3.3(10) | -14.8±1.7(18) |
| **100 nM PF05089771 (NaV1.7-selective inhibitor)** |  |  |  |  |
| Activation |  |  |  |  |
| $V_{1/2}$ | 4.0±1.2(12) | 9.0±1.4(12) | -16.8±0.8(9) | -14.9±1.3(11)<sup>b</sup> |
| $k$ | 9.5±1.0(12) | 9.4±1.5(12) | 4.5±0.7(9) | 8.0±1.3(11) |
| Inactivation |  |  |  |  |
| $V_{1/2}$ | -37.2±1.6(12) | -24.8±2.2(12)<sup>c</sup> | -33.3±1.0(9) | -35.7±2.4(11)<sup>c</sup> |
| $k$ | -10.9±1.5(12) | -11.4±1.9(12) | -8.2±1.0(9) | -13.4±2.3(11) |
| **DMSO (Control)** |  |  |  |  |
| Activation |  |  |  |  |
| $V_{1/2}$ | -22.5±4.1(14) | -21.26±2.6(26) | -16.1±1.8(26) | -12.7±2.8(19) |
| $k$ | 5.6±2.0(14) | 5.8±1.7(26) | 7.4±1.8(26) | 7.7±1.7(19) |
| Inactivation |  |  |  |  |
| $V_{1/2}$ | -58.2±3.1(14) | -46.1±1.4(26) | -45.1±2.0(26) | -47.0±1.7(19) |
| $k$ | -14.6±1.1(14) | -16.3±1.9(13) | -24.5 ±3.3(26) | -16.0±1.4(19) |
| **20 μM Pitstop2 (inhibitor of clathrin assembly)** |  |  |  |  |
| Activation |  |  |  |  |
| $V_{1/2}$ | -23.4±1.2(14) | -23.1±1.4(26) | -13.6±2.8(26) | -14.7±1.8(19) |
| $k$ | 7.6±1.1(14) | 4.9±1.8(26) | 7.9±0.7(26) | 7.7±1.1(19) |
| Inactivation |  |  |  |  |
| $V_{1/2}$ | -46.1±2.6(14) | -44.0±3.2(26) | -39.8±2.0(26) | -33.1±1.8(19) |
| $k$ | -21.3±3.5(14) | -16.6±1.9(26) | -14.4±2.0(26) | -17.3±1.5(19) |

<sup>a</sup>Values are means ± S.E.M. calculated from fits of the data from the indicated number of individual cells (in parentheses) to the Boltzmann equation; $V_{1/2}$ midpoint potential (mV) for voltage-dependent activation or inactivation; $k$ , slope factor. These values pertain to **Figures 5, 7**. Only statistically significant differences are indicated within the table
<sup>b</sup> $p=0.0330$ comparing vs. WT control vs. CRMP2<sup>K374A/K374A</sup> + PF05089771 (one-way ANOVA with Tukey's post hoc test)
<sup>c</sup> $p<0.0001$ comparing vs. WT control vs. CRMP2<sup>K374A/K374A</sup> + PF05089771 (one-way ANOVA with Tukey's post hoc test)

**Table 2.** Statistical analyses of behavioral experiments.

| Figure panel | Assay | Statistical test; findings | Post-hoc analysis (adjusted p-values) | Number of subjects |
| --- | --- | --- | --- | --- |
| Figure 3B, C | Marble burying test | | Mann-Whitney test<br>Females: $p = 0.5456$<br>Males: $p = 0.0.6918$ | WT females: $n = 9$<br>CRMP2<sup>K374A/K374A</sup><br>females: $n = 8$<br>WT males: $n = 10$<br>CRMP2<sup>K374A/K374A</sup><br>males: $n = 11$ |
| Figure 3E, F | Nestlet shredding test | | Mann-Whitney test<br>Females: $p > 0.9999$<br>Males: $p = 0.4490$ | WT females: $n = 9$<br>CRMP2<sup>K374A/K374A</sup><br>females: $n = 8$<br>WT males: $n = 11$<br>CRMP2<sup>K374A/K374A</sup><br>males: $n = 11$ |
| Figure 3G | Tail suspension test | Kruskal Wallis test<br>$P = 0.2353$ | Dunn's multiple comparisons test<br>WT females vs. CRMP2<sup>K374A/K374A</sup><br>females: $p > 0.9999$<br>WT females vs. WT males: $p > 0.9999$<br>WT females vs. CRMP2<sup>K374A/K374A</sup><br>males: $p > 0.9999$<br>CRMP2<sup>K374A/K374A</sup><br>females vs. WT males: $p = 0.3071$<br>CRMP2<sup>K374A/K374A</sup><br>females vs. CRMP2<sup>K374A/K374A</sup><br>males: $p = 0.8159$<br>WT males vs. CRMP2<sup>K374A/K374A</sup><br>males: $p > 0.9999$ | WT females: $n = 15$<br>CRMP2<sup>K374A/K374A</sup><br>females: $n = 15$<br>WT males: $n = 16$<br>CRMP2<sup>K374A/K374A</sup><br>males: $n = 12$ |
| Figure 3H | Elevated plus maze | Kruskal Wallis test<br>$P = 0.0004$ | Dunn's multiple comparisons test<br>WT females vs. CRMP2<sup>K374A/K374A</sup><br>females: $p = 0.0009$<br>WT females vs. WT males: $p > 0.9999$<br>WT females vs. CRMP2<sup>K374A/K374A</sup><br>males: $p = 0.0308$<br>CRMP2<sup>K374A/K374A</sup><br>females vs. WT males: $p = 0.0107$<br>CRMP2<sup>K374A/K374A</sup><br>females vs. CRMP2<sup>K374A/K374A</sup><br>males: $p > 0.9999$<br>WT males vs. CRMP2<sup>K374A/K374A</sup><br>males: $p = 0.0293$ | WT females: $n = 15$<br>CRMP2<sup>K374A/K374A</sup><br>females: $n = 15$<br>WT males: $n = 14$<br>CRMP2<sup>K374A/K374A</sup><br>males: $n = 13$ |
| Figure 4A, B | Hot plate at 48°C | | Mann Whitney test<br>WT females vs. CRMP2<sup>K374A/K374A</sup><br>females: $p = 0.0015$ | WT females: $n = 12$<br>CRMP2<sup>K374A/K374A</sup><br>females: $n = 15$<br>WT males: $n = 20$ |
|  |  |  | WT males vs.<br>CRMP2<sup>K374A/K374A</sup><br>males: p = 0.1302 | CRMP2<sup>K374A/K374A</sup><br>males: n = 22 |
| Figure 4C, D | Hot plate at 52°C |  | Mann Whitney test<br>WT females vs.<br>CRMP2<sup>K374A/K374A</sup><br>females: p = 0.0234<br>WT males vs.<br>CRMP2<sup>K374A/K374A</sup><br>males: p = 0.0007 | WT females: n = 12<br>CRMP2<sup>K374A/K374A</sup><br>females: n = 15<br>WT males: n = 20<br>CRMP2<sup>K374A/K374A</sup><br>males: n = 22 |
| Figure 4E, F | Hot plate at 55°C |  | Mann Whitney test<br>WT females vs.<br>CRMP2<sup>K374A/K374A</sup><br>females: p = 0.9454<br>WT males vs.<br>CRMP2<sup>K374A/K374A</sup><br>males: p = 0.7627 | WT females: n = 12<br>CRMP2<sup>K374A/K374A</sup><br>females: n = 15<br>WT males: n = 20<br>CRMP2<sup>K374A/K374A</sup><br>males: n = 22 |
| Figure 4G, H | Tail flick at 52°C |  | Mann Whitney test<br>WT females vs.<br>CRMP2<sup>K374A/K374A</sup><br>females: p = 0.7872<br>WT males vs.<br>CRMP2<sup>K374A/K374A</sup><br>males: p = 0.3605 | WT females: n = 15<br>CRMP2<sup>K374A/K374A</sup><br>females: n = 17<br>WT males: n = 14<br>CRMP2<sup>K374A/K374A</sup><br>males: n = 17 |
| Figure 9 | Veratridine induced<br>paw licking |  | Mann Whitney test<br>Phase 1 –<br>WT females vs.<br>CRMP2<sup>K374A/K374A</sup><br>females: p = 0.0115<br>WT males vs.<br>CRMP2<sup>K374A/K374A</sup><br>males: p = 0.7925<br>Phase 2 –<br>WT females vs.<br>CRMP2<sup>K374A/K374A</sup><br>females: p = 0.1656<br>WT males vs.<br>CRMP2<sup>K374A/K374A</sup><br>males: p = 0.01641 | WT females: n = 6<br>CRMP2<sup>K374A/K374A</sup><br>females: n = 8<br>WT males: n = 7<br>CRMP2<sup>K374A/K374A</sup><br>males: n = 8 |
| Figure 10A | Spared nerve injury | Two-way ANOVA with<br>Sidak's post hoc test | Females WT vs<br>CRMP2<sup>K374A/K374A</sup>;<br>Baseline p>0.9999<br>7 days p=0.0371<br>14 days p=0.0379<br>21 days p=0.0066<br>28 days p=0.0018<br>35 days p<0.0001 | WT females: n=13<br>CRMP2<sup>K374A/K374A</sup><br>females: n=13 |
| Figure 10B | Spared nerve injury |  | Mann Whitney test<br>Females WT vs<br>CRMP2<sup>K374A/K374A</sup>;<br>p=0.0005 | WT females: n=13<br>CRMP2<sup>K374A/K374A</sup><br>females: n=13 |
| Figure 10C | Spared nerve injury | Two-way ANOVA with<br>Sidak's post hoc test | Males WT vs<br>CRMP2<sup>K374A/K374A</sup>;<br>Baseline p>0.9999<br>7 days p=0.0316<br>14 days p=0.0004<br>21 days p=0.0022<br>28 days p<0.0001<br>35 days p<0.0001 | WT males: n=12<br>CRMP2<sup>K374A/K374A</sup><br>males: n=12 |
| Figure 10D | Spared nerve injury | Mann Whitney test | Mann Whitney test | WT males: n=12<br>CRMP2<sup>K374A/K374A</sup><br>males: n=12 |
|  |  |  | Males WT vs<br>CRMP2<sup>K374A/K374A</sup>;<br>p<0.0001 |  |
| Figure 11B, D | Formalin induced<br>licking |  | Mann Whitney test<br>WT females vs.<br>CRMP2<sup>K374A/K374A</sup><br>females: p = 0.1659<br>WT males vs.<br>CRMP2<sup>K374A/K374A</sup><br>males: p = 0.8317 | WT females: n = 9<br>CRMP2<sup>K374A/K374A</sup><br>females: n = 7<br>WT males: n = 6<br>CRMP2<sup>K374A/K374A</sup><br>males: n = 10 |
| Figure 11E, F | Acetic acid induced<br>writting | Kruskal Wallis test | Dunn's multiple<br>comparisons test<br>WT females vs.<br>CRMP2<sup>K374A/K374A</sup><br>females: p >0.9999<br>WT females vs. WT<br>males: p = 0.9298<br>WT females vs.<br>CRMP2<sup>K374A/K374A</sup><br>males: p >0.9999<br>CRMP2<sup>K374A/K374A</sup><br>females vs. WT males:<br>p = 0.6262<br>CRMP2<sup>K374A/K374A</sup><br>females vs.<br>CRMP2<sup>K374A/K374A</sup><br>males: p = 0.8279<br>WT males vs.<br>CRMP2<sup>K374A/K374A</sup><br>males: p >0.9999 | WT females: n = 9<br>CRMP2<sup>K374A/K374A</sup><br>females: n = 9<br>WT males: n = 9<br>CRMP2<sup>K374A/K374A</sup><br>males: n = 9 |
| Figure 11G | Paw incision – Von<br>Frey mechanical<br>thresholds | Two-way ANOVA with<br>Sidak's post hoc test | Females WT vs<br>CRMP2<sup>K374A/K374A</sup>;<br>Baseline p=0.7210<br>24 hours p=0.9122<br>After GBP p=0.9890 | WT females: n=9<br>CRMP2<sup>K374A/K374A</sup><br>females: n=9 |
| Figure 11H | Paw incision – Von<br>Frey mechanical<br>thresholds | Two-way ANOVA with<br>Sidak's post hoc test | Males WT vs<br>CRMP2<sup>K374A/K374A</sup>;<br>Baseline p=0.4853<br>24 hours p=0.8050<br>After GBP p=0.5181 | WT males: n=9<br>CRMP2<sup>K374A/K374A</sup><br>males: n=9 |

## 3. Results

### 3.1. CRMP2 SUMOylation is abolished in CRMP2^K374A/K374A^ mice

Since the Lys374 residue of CRMP2 is the major SUMOylation site [17-19], we generated CRMP2 SUMO-null knock-in (CRMP2^K374A/K374A^) mice in which the lysine residue was replaced with alanine (**Figure 1A, B**). To verify that CRMP2 SUMOylation was impaired by the CRMP2^K374A/K374A^ mutation in situ, we used the proximity ligation assay (PLA) to detect SUMOylated CRMP2 in DRG neurons from either WT or CRMP2^K374A/K374A^ mice. PLA allows for a fluorescent punctum to be detected at the site(s) where two proteins are within 30-nm or less of each other, thereby permitting detection of protein-complex formation at endogenous levels. In situ PLA utilizes antibodies to which DNA oligonucleotides have been attached as probes. Proximal binding of the PLA probes will allow for the hybridization of two additional oligonucleotides to the PLA probes. These oligonucleotides can then be ligated to form a circular DNA reporter molecule, which is amplified by rolling cycle amplification. The resulting threads of single stranded DNA will collapse into a bundle – the rolling cycle amplification product. These products are detected by fluorescence labeled oligonucleotides to generate individual bright fluorescent signals in the place where complex formation between the protein antigens was detected. We stained DRG neurons using a CRMP2 and a SUMO1 antibody followed by the PLA protocol (**Figure 1C**). The number of PLA puncta was then quantified per DRG neuron. We found no difference in the number of SUMOylated CRMP2 puncta per neuron between WT male (4.08 ± 0.74, n=12) and female (4.43 ± 0.55, n=14) mice (**Figure 1D**). In contrast, in DRGs from CRMP2^K374A/K374A^ female and male mice, the SUMOylated CRMP2 PLA signals were completely suppressed. No signals were detected in absence of either of the antibodies (not shown). These observations indicated that the CRMP2^K374A/K374A^ knock-in mutation produced a loss of Lys374 SUMOylation of CRMP2 in DRG neurons.

### 3.2. Long term potentiation (LTP) is not affected in CRMP2^K374A/K374A^ mice

CRMP2 is known to be expressed in the hippocampus and is involved in determining dendritic patterning in this brain region [34]. In CRMP2 knockout mice, long-term potentiation (LTP) is impaired and associated with poorer performance in behavioral tests assessing learning [74]. To investigate whether LTP was impacted in CRMP2^K374A/K374A^ mice, we performed electrophysiological recordings of the Schaffer collateral CA1 circuit in coronal hippocampal brain slices acutely prepared from wildtype and CRMP2^K374A/K374A^ mice using microelectrode arrays. First, we examined the input output relationship to determine if there were differences in basal excitability levels between WT and CRMP2^K374A/K374A^ groups in both male and female animals and found no significant differences between the groups (**Figure 2A, E**). Next, to test if the K374A mutation caused changes in presynaptic plasticity, we measured the paired pulse ratio with pulse intervals of 15, 50, 150 and 400-ms. We found that both sexes demonstrated no difference in presynaptic plasticity as measured by their paired pulse ratios (**Figure 2B, F**). We next utilized a theta burst stimulation protocol to induce LTP, which allowed us to determine if differences in postsynaptic plasticity existed in either group (**Figure 2C, G**) [39]. We assessed the level of potentiation in each group during the final 5 minutes of the 60-minute post-stimulation recording period to quantify differences in LTP between groups. There were no significant differences between groups in the level of potentiation observed (**Figure 2D, H**). Taken together, these results suggest that the CRMP2^K374A/K374A^ genotype does not impact basal excitability, presynaptic or postsynaptic hippocampal plasticity in the hippocampal CA1 circuit. Our findings suggest that targeting Lys374 of CRMP2 does not produce significant changes to the innate physiological properties of neural circuitry involved in learning and memory behavior. Furthermore, as the hippocampus is the primary neuronal substrate of short-term memory and spatial orientation, the lack of differences between WT and CRMP2^K374A/K374A^ groups in our experiments suggests that these capacities remain intact in these knock-in animals.

**Figure 2.**
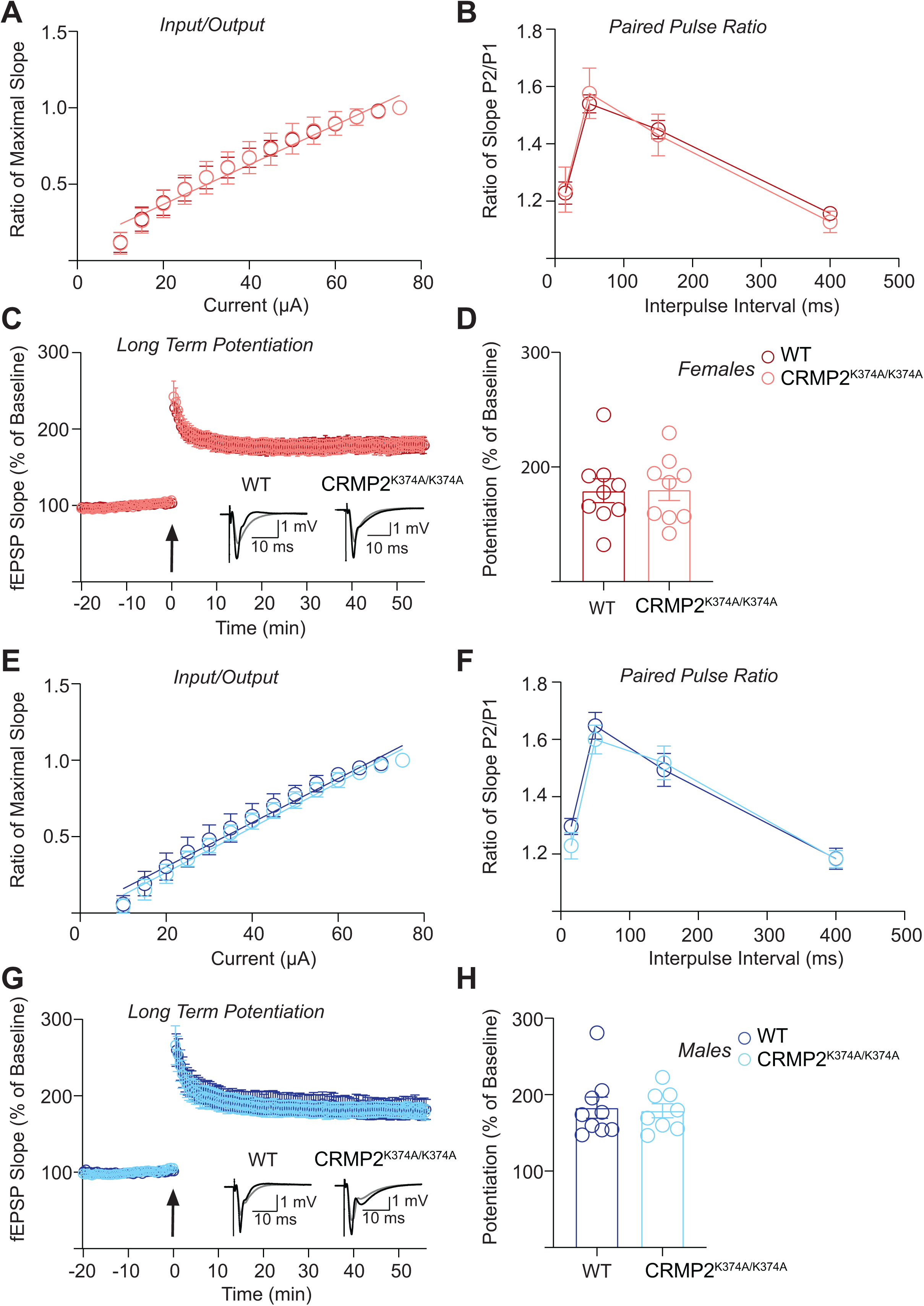
Basal excitability, presynaptic plasticity, and long-term potentiation in hippocampal neurons are not changed between wildtype and CRMP2^K374A/K374A^ knock-in mice. (**A**) Input output relationship of K374A wild type (WT) and homozygous (CRMP2^K374A/K374A^) groups obtained from female mice. Data was fit using nonlinear regression and a linear approximation of the slope that best fit the data from each group was obtained. Analysis of covariance was used to determine whether the slope of each group was significantly different. The line of best fit for both groups has the same slope [P = 0.9699, F(2, 290) = 0.03061]. WT N = 9 slices (3 animals), CRMP2^K374A/K374A^ N = 12 slices (4 animals). (**B**) Paired pulse facilitation ratio obtained from K374A WT and CRMP2^K374A/K374A^ female mice to assess changes in presynaptic plasticity. Pulse intervals were 15, 50, 150 and 400 ms. Two-way ANOVA was performed to identify significant differences between groups, and none were found [P = 0.9939, F(1,76) = 5.885 × 10^−5^]. WT N = 9 slices (3 animals), CRMP2^K374A/K374A^ N = 12 slices (4 animals). (**C**) Long term potentiation of K374A WT and CRMP2^K374A/K374A^ female mice represented as a percentage of averaged 20-minute baseline response. Arrow indicates time 0 when the theta burst stimulation (TBS) protocol was applied to the stimulation electrode positioned in the Schaffer Collateral projection from CA3 to CA1 region of the hippocampus. Inset shows representative fEPSP traces before (Gray) and after (Black) TBS application for both WT and CRMP2^K374A/K374A^ groups. WT N = 9 slices (3 animals), CRMP2^K374A/K374A^ N = 9 slices (4 animals). (**D**) Averaged level of potentiation during the last 5 minutes of the recording period for comparison between WT and CRMP2^K374A/K374A^ groups in females. Averaged potentiation was compared between groups using a two-tailed unpaired t-test, which did not reach significance [P = 0.9431, t = 0.07245, df = 16]. WT N = 9 slices (3 animals), CRMP2^K374A/K374A^ N = 9 slices (4 animals). (**E**) Input output relationship of WT and CRMP2^K374A/K374A^ obtained from male mice. Data was fit using nonlinear regression and a linear approximation of the slope that best fit the data from each group was obtained. Analysis of covariance was used to determine whether the slope of each group was significantly different. The line of best fit for both groups has the same slope [P = 0.5191, F(1,276) = 0.4167]. WT N = 11 slices (3 animals), CRMP2^K374A/K374A^ N = 9 slices (3 animals). (**F**) Paired pulse facilitation ratio obtained from WT and CRMP2^K374A/K374A^ male mice to assess changes in presynaptic plasticity. Pulse intervals were 15, 50, 150 and 400 ms. Two-way ANOVA was performed to identify significant differences between groups, and none were found [P = 0.4716, F(1,72) = 0.5238]. WT N = 11 slices (3 animals), CRMP2^K374A/K374A^ N = 9 slices (3 animals). (**G**) Long term potentiation of WT and CRMP2^K374A/K374A^ male mice represented as a percentage of averaged 20-minute baseline response. Arrow indicates time 0 when the theta burst stimulation (TBS) protocol was applied to the stimulation electrode positioned in the Schaffer Collateral projection from CA3 to CA1 region of the hippocampus. Inset shows representative fEPSP traces before (Gray) and after (Black) TBS application for both WT and CRMP2^K374A/K374A^ groups. WT N = 9 slices (3 animals), CRMP2^K374A/K374A^ N = 8 slices (3 animals). (**H**) Averaged level of potentiation during the last 5 minutes of the recording period for comparison between WT and CRMP2^K374A/K374A^ groups in males. Averaged potentiation was compared between groups using a two-tailed unpaired t-test, which did not reach significance [P = 0.8167, t = 0.2359, df = 15]. WT N = 9 slices (3 animals), CRMP2^K374A/K374A^ N = 8 slices (3 animals). The experiments were conducted by investigators blinded to the genotype. Error bars indicate mean ± SEM.

### 3.3. Evaluation of cognitive behaviors in CRMP2^K374A/K374A^ mice

A key role for CRMP2 in cognitive processing was inferred from a previous study using global CRMP2 knockout mice [55]. Thus, we asked if the CRMP2^K374A/K374A^ knock-in mice might present with abnormal behaviors related to repetitive, compulsive-like behaviors. This was initially evaluated using the marble burying test (**Figure 3A**); increased marble burying is associated with obsessive-compulsive disorder (OCD) behaviors [3]. Here, we observed no effect of sex or genotype (**Figure 3A-F**). We also performed the nestlet shredding test where increased shredding is associated with anxiety and OCD [57]. Again, there was no difference in nestlet shredding behavior across sexes and genotypes. Together, these observations indicated that the CRMP2^K374A/K374A^ mutation is not related to OCD-like behaviors. Using the tail suspension test [13], we found no evidence of altered depression-like behaviors in female or male CRMP2^K374A/K374A^ mice (**Figure 3G**). Finally, because CRMP2 knockout mice exhibited anxiolytic behaviors [55], we used the elevated plus maze (EPM) to test for anxiety. CRMP2^K374A/K374A^ mice spent more time in the open arms of the EPM (**Figure 3H**). These results show that the CRMP2^K374A/K374A^ mutation had no effect on OCD and depression-like behaviors but conferred an anxiolytic phenotype on the knock-in mice.

**Figure 3.**
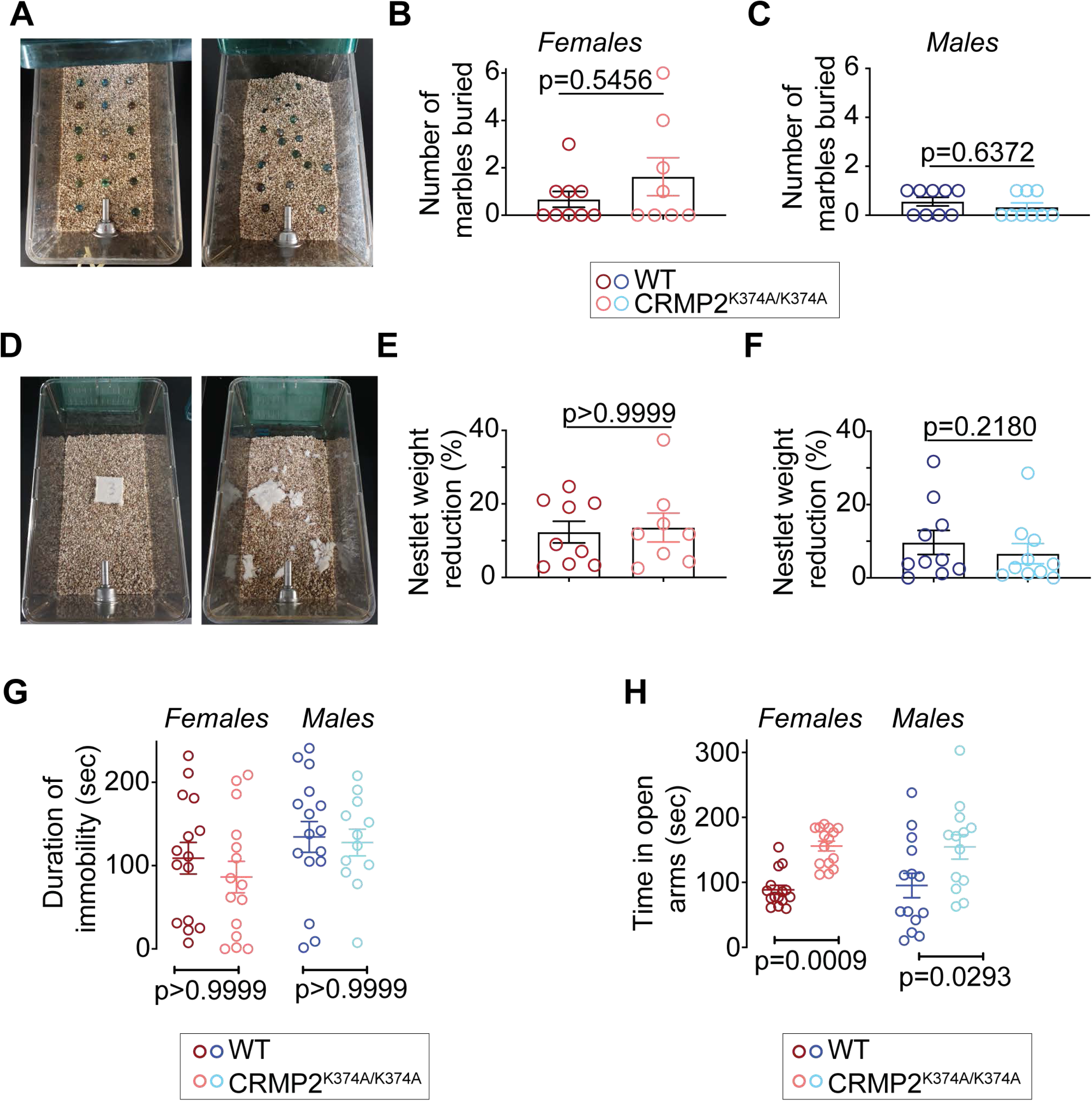
CRMP2^K374A/K374A^ knock-in mice do not exhibit compulsive-like or depression-associated but have reduced anxiety-associated behaviors. (**A**) Each cage was filled with 5cm of bedding chips lightly pressed to create a flat, even surface. 17 glass marbles (1.5cm in diameter) were distributed on the bedding surface. Mice were placed in the cages (1 mouse/cage) and allowed to behave freely for 30 minutes. After 30 minutes the number of marbles at least at least 2/3^rds^ buried/covered with bedding were counted. No differences between genotypes was observed in females (**B**) or males (**C**). (**D**) For the nestlet test of compulsive-like behaviors, a nestlet was weighed and placed in the center of the cage. Mice were placed in the cages (1 mouse/cage) and allowed to behave freely for 30 minutes. Afterward, the largest remaining piece of nestlet was weighed. No differences between genotypes were observed in females (**E**) or males (**F**). (**G**) Immobility time in the tail suspension test was no different between sexes or genotypes. (**H**) Time spent in the open arms of the elevated plus-maze was reduced in the female and male CRMP2^K374A/K374A^ mice compared to their WT counterparts. See statistical analysis described in **Table 2**. Error bars indicate mean ± SEM.

### 3.4. Evaluation of the consequences of the CRMP2^K374A/K374A^ genotype on the immune cell landscape

Although not well understood, CRMP2 was reported to be involved in T lymphocyte migration and recruitment of immune cells into the brain [68; 69]. Therefore, we next asked if the CRMP2^K374A/K374A^ mutation could impact immune cell populations. We performed fluorescence activated cell sorting (FACS) analyses of immune cells from brain, spinal cord, blood and spleen of naïve male and female wildtype and CRMP2^K374A/K374A^ mice. In brain and spinal cord, we found no difference in the abundance of endothelial cells, astrocytes, oligodendrocytes, neurons or microglia (**Supplementary Figure 2, Supplementary Table 1**). In blood, we found an increased proportion of CD8+ T lymphocytes in male CRMP2^K374A/K374A^ mice compared to wildtype mice, whereas other cell populations such as helper (CD4+), regulatory (CD25+) or activated (CD69+) T cells remained unchanged in blood and spleen from either genotype (**Supplementary Figure 3, Supplementary Table 1**). This broad characterization of the immune cell landscape in CRMP2^K374A/K374A^ mice shows that the K374A mutation had negligible effects on the immune system.

### 3.5. CRMP2^K374A/K374A^ mice have thermal analgesia

The mice’s responses to acute noxious thermal stimuli was examined using the hot-plate (measures supraspinal responses to nociception) and tail-flick (reflexive nociception response) tests. Compared with wildtype mice, female CRMP2^K374A/K374A^ mice displayed increased response latency to 48°C and 52°C stimuli but retained equal sensitivity to a 55°C stimulus (**Figure 4A, C and E**). In contrast, male CRMP2^K374A/K374A^ mice exhibited a higher latency to respond only to a 52°C nociceptive stimulus compared to wildtype mice (**Figure 4B, D, F**). Next, we used the tail flick test to measure a response that would be independent of higher brain functions. In both female and male mice of either genotype no altered response latency was detected in our tail flick experiment (52°C) (**Figure 4G-H**). Together. these results highlight a role for CRMP2 SUMOylation in regulating thermal pain sensation.

**Figure 4.**
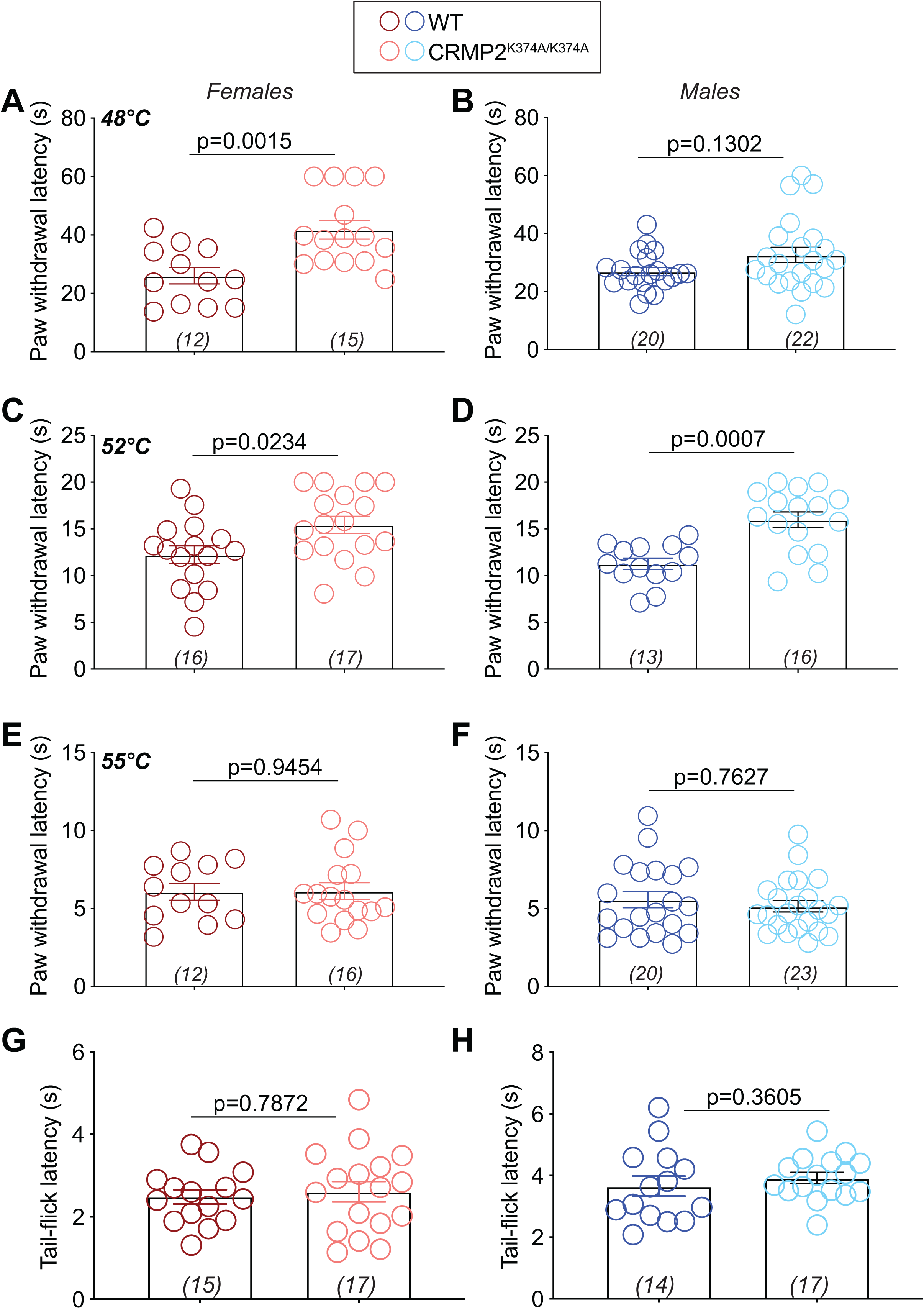
Preventing CRMP2 SUMOylation decreases thermal sensitivity in males and females. Latencies to respond to noxious heat (48°C, 52°C, or 55°C) in the hot plate (**A**-**F**) or tail-flick (52°C) (**G, H**) tests in female and male wildtype (WT) and CRMP2^K374A/K374A^ mice. Paw withdrawal latency was increased in heterozygous males at 48°C vs. wildtype males and increased at 52°C between wildtype and homozygotes males. There were no changes in latencies at the highest temperature. At 48°C, female CRMP2^K374A/K374A^ mice exhibited a higher latency to respond to the nociceptive stimulus compared to wildtype mice. There were no changes in latencies at 52°C or 55°C. There were no changes in the hot-plate test under any of the conditions; the test was stopped at the cutoff time of 10-s. See statistical analysis described in **Table 2**. Error bars indicate mean ± SEM.

### 3.6. NaV1.7 currents are decreased in female CRMP2^K374A/K374A^ mice

We previously reported that CRMP2 expression and CRMP2 SUMOylation regulate NaV1.7 function [17; 18; 22]. To test if this regulation was affected in our knock-in mice, we measured sodium currents in DRG neurons isolated from WT or CRMP2^K374A/K374A^ male and female mice (**Figure 5A**). In small diameter DRGs from female mice, the CRMP2^K374A/K374A^ genotype was associated with an ∼40% decrease in total sodium current compared to DRGs from WT mice (**Figure 5B**), thus corroborating our previous observations. Surprisingly, in male mice, we found no difference in total sodium currents between genotypes (**Figure 5B**). These observations suggested a sex-specific regulation of sodium currents imposed by the CRMP2^K374A/K374A^ mutation. To assess the contribution of NaV1.7 to the total sodium currents in our mice, we used the NaV1.7-specific blocker PF05089771 [1; 42]. Here, ∼76% of the sodium current was PF05089771-sensitive (i.e., carried via NaV1.7) in female DRGs from both WT and CRMP2^K374A/K374A^ mice (**Figure 5C-D**). Voltage-dependence of activation and inactivation were both shifted to the right (i.e. depolarized) by PF05089771 in DRGs from female WT and CRMP2^K374A/K374A^ mice (**Figure 5E, F**). Voltage-dependence of activation remained unchanged across genotypes and treatment conditions in male DRGs. However, the voltage-dependence of inactivation was shifted to the right by 18.4 mV in the presence of PF05089771 (**Figure 5I, J**). In DRGs from male mice, adding PF05089771 lead to a decrease of total sodium currents by 53% and 58% from WT and CRMP2^K374A/K374A^ genotypes, respectively (**Figure 5G-H**).

**Figure 5.**
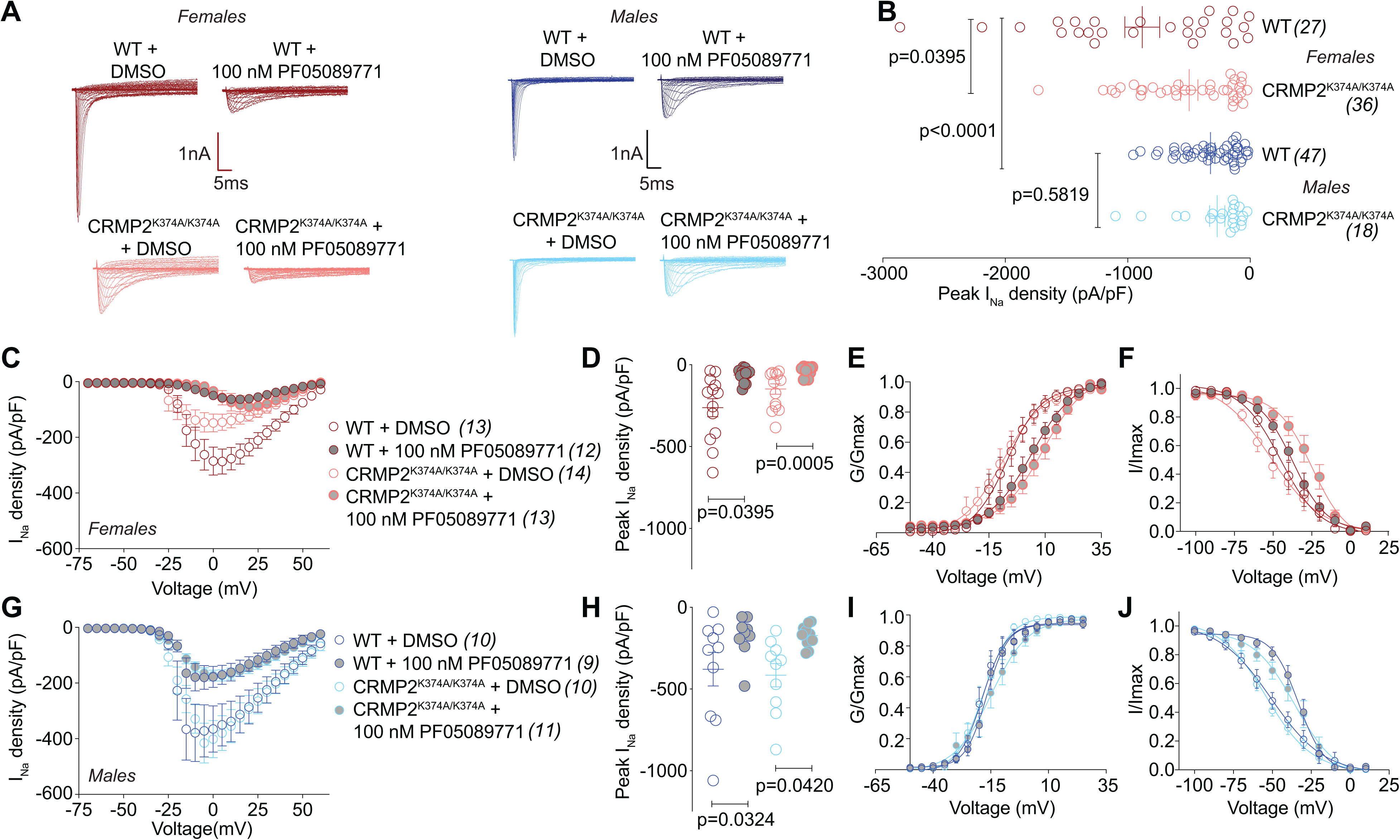
Total sodium currents and PF-05089771-sensitive NaV1.7 currents are reduced in female, but not male, DRGs from CRMP2^K374A/K374A^ knock-in mice. (**A**) Representative current traces recorded from small-sized DRGs neurons isolated from female and male wildtype (WT) and CRMP2^K374A/K374A^ knock-in mice in response to depolarization steps from −70 to +60 mV from a holding potential of −60 mV. (**B**) Summary of normalized peak currents (in picoamperes/picofarads, pA/pF) from DRG neurons cultured from female and male wildtype (WT) and CRMP2^K374A/K374A^ knock-in mice. Total sodium current was similar between wildtype and homozygous male mice (n=18-47 cells/condition, p = 0.5819, Kruskal-Wallis test with Dunn’s post hoc). NaV1.7 currents were significantly smaller in DRGs from homozygous female mice vs. DRG neurons from wildtype female (n=27-36 cells/condition, p = 0.03959, Kruskal-Wallis test with Dunn’s post hoc) or the both genotypes of male DRGs (p <0.0001, Kruskal-Wallis test with Dunn’s post hoc). The NaV1.7-selective inhibitor PF-05089771 [1] (100 nM, 5-15 min) was used to assess the extent of sodium currents carried by NaV1.7 in DRGs neurons isolated from female and male WT and CRMP2^K374A/K374A^ knock-in mice. Summary of current-voltage curves (**C** – *females*; **G** – *males*) and normalized peak (**D** – *females*; **H** – *males*) currents (pA/pF) from DRG neurons of WT and CRMP2^K374A/K374A^ knock-in mice. Boltzmann fits for normalized conductance G/G_max_ voltage relations for voltage dependent activation (**E, I**) and inactivation (**F, J**) of the sensory neurons as indicated. Half-maximal activation and inactivation (V_1/2_) and slope values (*k*) for activation and inactivation are presented in **Table 1**. Error bars indicate mean ± SEM.

We interrogated our cell culture conditions to investigate the reason underlying decreased NaV1.7 currents in female CRMP2^K374A/K374A^ mice. In vivo, multiple growth factors are present in the DRG micro-environment, lack of these factors in vitro may result in a down-regulation of either NaV1.7 or CRMP2. Because of their role in neuropathic pain and in regulating CRMP2 [30; 54], we chose to test if nerve growth factor (NGF, 125 ng/ml) or brain-derived neurotrophic factor (BDNF, 20ng/ml) could ‘normalize’ NaV1.7 currents in cultured DRG neurons from male CRMP2^K374A/K374A^ mice. Adding either NGF or BDNF to DRGs did not affect NaV1.7 currents compared to untreated DRGs from WT or CRMP2^K374A/K374A^ mice (**Supplementary Figure 4**). Together, these results show that the K374A knock-in mutation decreased total sodium current by inhibiting NaV1.7 in DRGs from female, but not male, mice. We did not observe any changes in voltage-gated calcium or potassium currents in DRGs isolated from female and male mice of either genotype (**Supplementary Figures 5-7**). This finding underscores the specificity of CRMP2 SUMOylation in the regulation of NaV1.7.

### 3.7. NaV1.7 trafficking and NaV1.7–CRMP2 interaction are reduced in DRGs from female CRMP2^K374A/K374A^ mice

Our previous work established that: (*i*) CRMP2 SUMOylation regulates the membrane localization of NaV1.7 [17; 19], and (ii) when deSUMOylated, CRMP2 binding to NaV1.7 is lost [17]. Our data above shows that NaV1.7 currents are decreased only in female DRGs, but whether NaV1.7 surface expression and CRMP2-NaV1.7 interactions are also sexually dimorphic is not known. Consequently, we first tested if the interaction between CRMP2 and NaV1.7 was affected equally in DRGs from both sexes of CRMP2^K374A/K374A^ mice. In DRG neurons, we stained for NaV1.7 and CRMP2 prior to developing the PLA signal in DRG neurons (**Figure 6A**). We quantified the number of puncta per neuron and found significantly less CRMP2-NaV1.7 PLA signal in DRGs from WT male mice compared to DRGs from WT female mice (**Figure 6B**). Equally importantly, we found that the CRMP2^K374A/K374A^ mutation significantly decreased the CRMP2/NaV1.7 PLA in DRGs from female, but not male, mice. Next, to test if this loss of CRMP2-NaV1.7 binding was related to altered membrane localization of NaV1.7, we stained DRG neurons for NaV1.7 and analyzed the amount of surface NaV1.7 using confocal microscopy (**Figure 6C**). In DRGs from female CRMP2^K374A/K374A^ mice, the membrane localization was decreased by ∼50% compared to DRGS from WT mice (**Figure 6D**). In DRGs from male mice, although we observed a more heterogenous distribution of NaV1.7 membrane localization in wildtype mice (**Figure 6D**), but the CRMP2^K374A/K374A^ mutation did not significantly decrease membrane localized NaV1.7 in male mice (**Figure 6D**).

**Figure 6.**
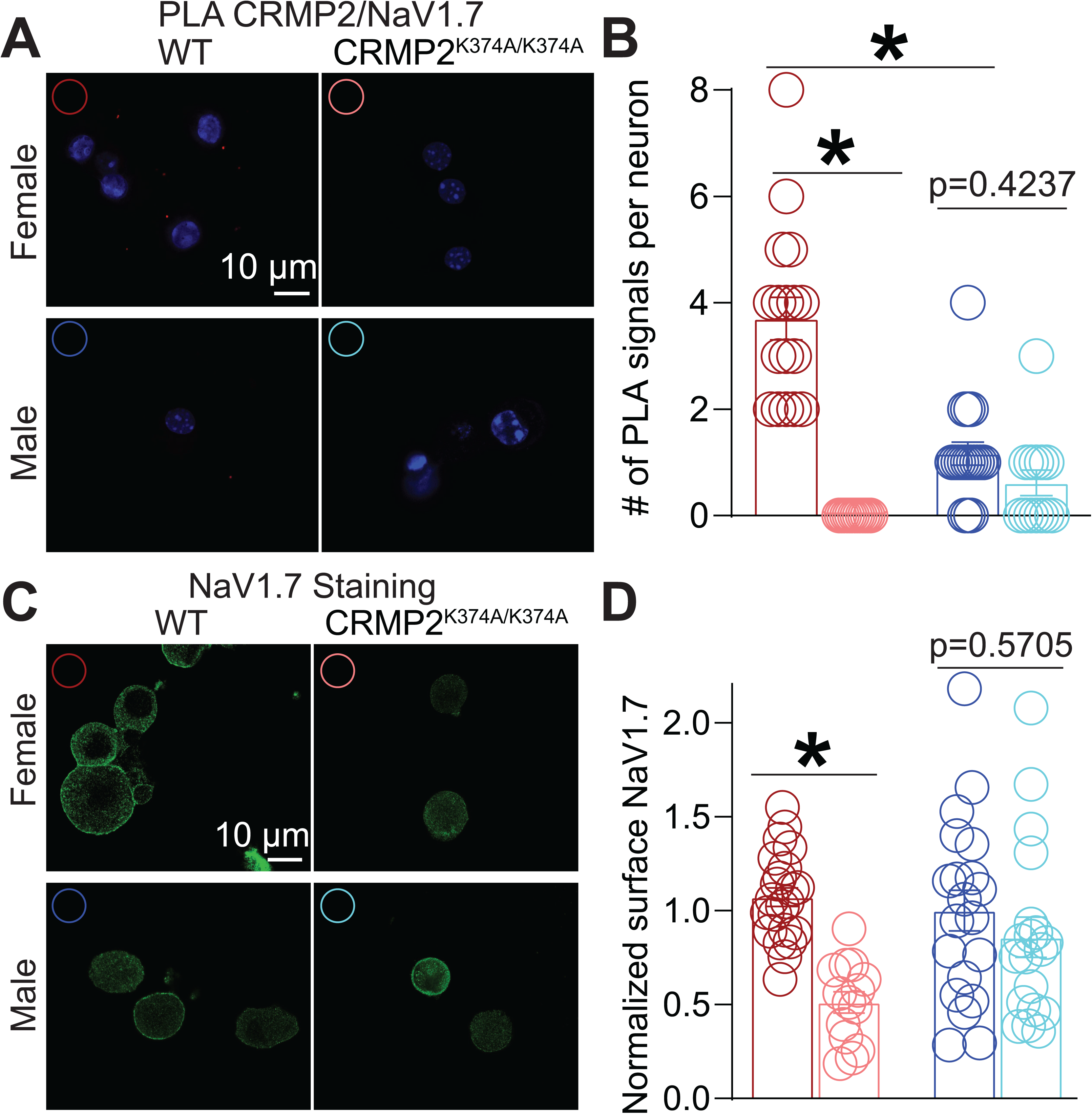
Surface NaV1.7 expression and binding of NaV1.7 to CRMP2 are reduced in female, but not male, DRGs from CRMP2^K374A/K374A^ knock-in mice. (**A**) Representative images of mouse DRG cultures following proximity ligation assay (PLA) between CRMP2 and NaV1.7. The PLA immunofluorescence labeled sites of interaction between CRMP2 and NaV1.7 (red puncta). Additionally, nuclei are labeled with the nuclear labeling dye 4’,6-diamidino-2-phenylindole (DAPI). Scale bar: 10 µm. (**B**) Quantification of PLA puncta per neuron shows that in DRGs from CRMP2^K374A/K374A^ mice, the number of NaV1.7-CRMP2 interactions was significantly reduced compared to wildtype DRGs from female mice (Kruskal-Wallis test with Dunn’s multiple comparison post-hoc test: p<0.0001 comparing female WT vs. female CRMP2^K374A/K374A^; p=0.4237 comparing male WT vs. male CRMP2^K374A/K374A^; and p=0.0016 comparing male WT vs. female WT; n=13-18 cells). (**C**) Representative confocal images of mouse DRG cultures labeled with an antibody against NaV1.7. (**D**) Quantification of normalized surface expression of NaV1.7 per neuron shows that in DRGs from CRMP2^K374A/K374A^ mice, the surface expression of NaV1.7 was significantly reduced compared to wildtype DRGs from female mice, whereas in male mouse DRGs there was no difference in expression between the genotypes (Kruskal-Wallis test with Dunn’s multiple comparison post-hoc test: p<0.0001 comparing female WT vs. female CRMP2^K374A/K374A^; p=0.5705 comparing male WT vs. male CRMP2^K374A/K374A^; and p=0.8994 comparing male WT vs. female WT; n=14-22 cells). Error bars indicate mean ± SEM.

As a corollary to these findings we tested the functional consequence of reduced CRMP2-NaV1.7 binding and reduced NaV1.7 surface expression. We had previously shown that loss of CRMP2 SUMOylation prevents NaV1.7 membrane localization by inducing its endocytosis. This was inferred from findings with Pitstop2, a clathrin assembly inhibitor [70], which prevented current density reductions imposed by loss of CRMP2 SUMOylation [17]. Inhibiting clathrin assembly with Pitstop2 negated the inhibition of sodium currents observed in DRGs from female CRMP2^K374A/K374A^ mice. In other words, application of Pitstop2 restored the decreased sodium currents back to the levels observed in DRGs from WT female mice (**Figure 7A, B**). No change in the biophysical properties of activation or inactivation was observed in control- or Pitstop2-treated samples from either genotype (**Figure 7C, D**). In contrast, in DRGs from male mice, inhibiting clathrin assembly had no effect (**Figure 7E, F**), thus confirming our observations that NaV1.7 trafficking was not affected by the CRMP2^K374A/K374A^ genotype in male mice (**Figure 6**). As before, no change of biophysical properties was observed in control- or Pitstop2-treated DRGs from male mice of either genotype (**Figure 7G, H**). Altogether, our results uncover a previously unknown sex difference in CRMP2-dependent regulation of NaV1.7 trafficking and function in DRG neurons.

**Figure 7.**
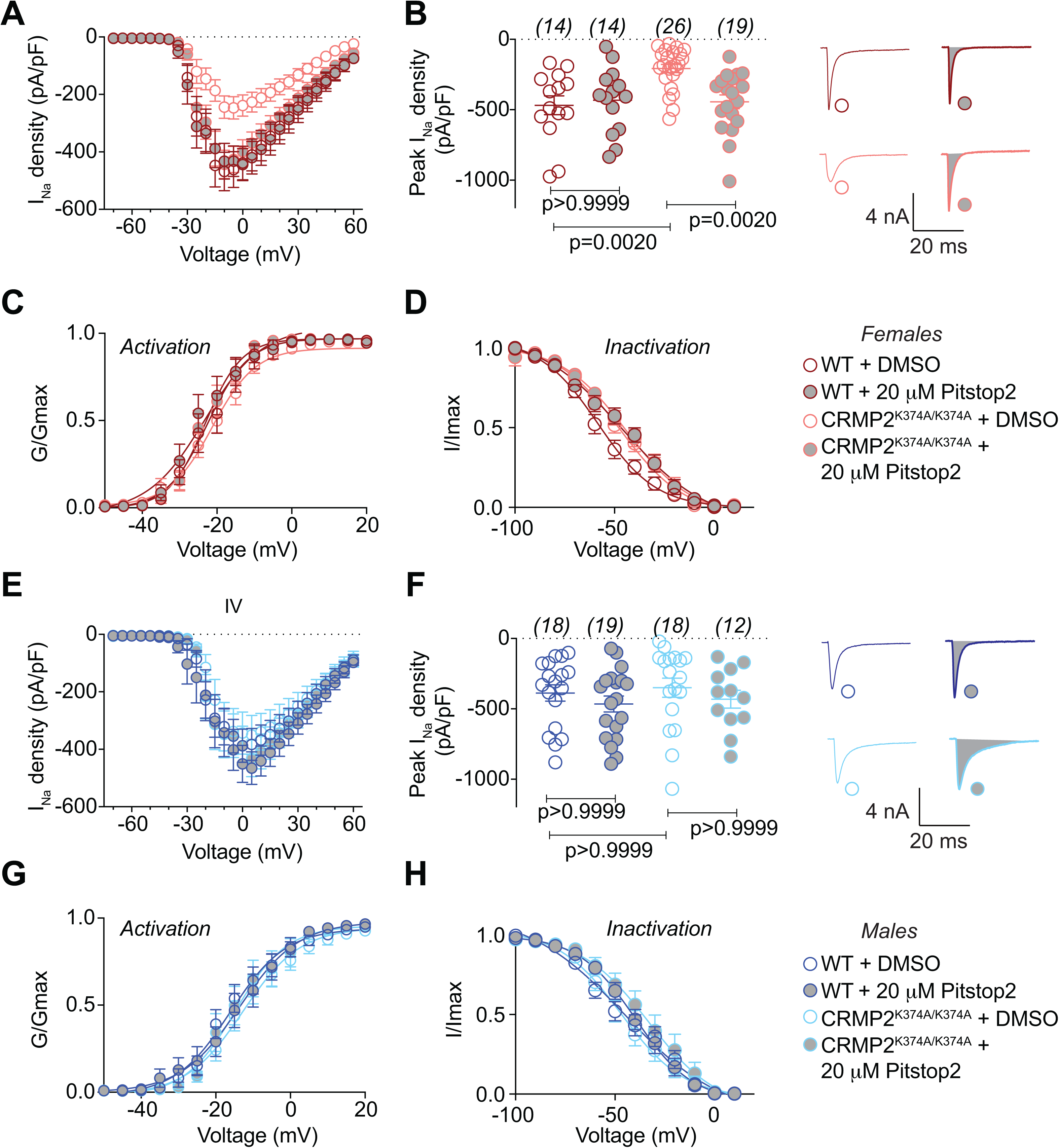
Reduction of NaV1.7 currents in DRGs from female but not male CRMP2^K374A/K374A^ knock-in mice are normalized by inhibition of clathrin coated endocytosis with Pitstop2. Summary of current-voltage curves (**A** – female, **E** – *male*) and normalized peak (**B** – female, **F** – *male*) currents (pA/pF) from small-to-medium diameter DRG neurons of WT and CRMP2^K374A/K374A^ knock-in mice (n = 14-26). In some experiments, currents were recorded following 30 minute application of 20 µM of Pitstop2, a clathrin-mediated endocytosis inhibitor [70]. Representative traces from female DRGs are displayed adjacent to the scatter graphs in **B**. Boltzmann fits for normalized conductance G/G_max_ voltage relations for voltage dependent activation (**C** – female, **G** – *male*) and inactivation (**D** – female, **H** – *male*) of the sensory neurons as indicated. Half-maximal activation and inactivation (V_1/2_) and slope values (*k*) for activation and inactivation are presented in **Table 1**. There were no significant differences in V_1/2_ and *k* values of activation and inactivation in either sex or genotype with and without treatment with Pitstop2 (One-way ANOVA with Tukey’s post hoc test). Error bars indicate mean ± SEM.

### 3.8. CRMP2^K374A/K374A^ has no impact on basal spinal spontaneous excitatory post-synaptic currents (sEPSCs)

We previously reported that CRMP2 phosphorylation by cyclin dependent kinase 5 (Cdk5) affected the frequency of sEPSCs in the spinal dorsal horn [73]. This function was hypothesized to be linked to CRMP2’s regulation of both the N-type voltage-gated calcium channel (CaV2.2) as well as NaV1.7. CaV2.2’s role in neurotransmission is to trigger the release of synaptic vesicles However, its activation relies on a prior depolarization event, triggered by opening of voltage gated sodium channels, particularly NaV1.7 in lamina I/II of the dorsal horn of the spinal cord [2]. Along with CaV2.2, NaV1.7 is another major determinant of synaptic transmission. Thus, we used slice electrophysiology to test if SUMOylated CRMP2 could impact spinal neurotransmission. Using whole-cell patch clamp, we recorded sEPSCs in neurons of the *Substantia Gelatinosa* (SG) region of the lumbar dorsal horn from CRMP2^K374A/K374A^ and wildtype mice of both genders. There was no change in amplitude or frequency of sEPSCs between CRMP2^K374A/K374A^ and wildtype mice of either sex (**Figure 8**).

**Figure 8.**
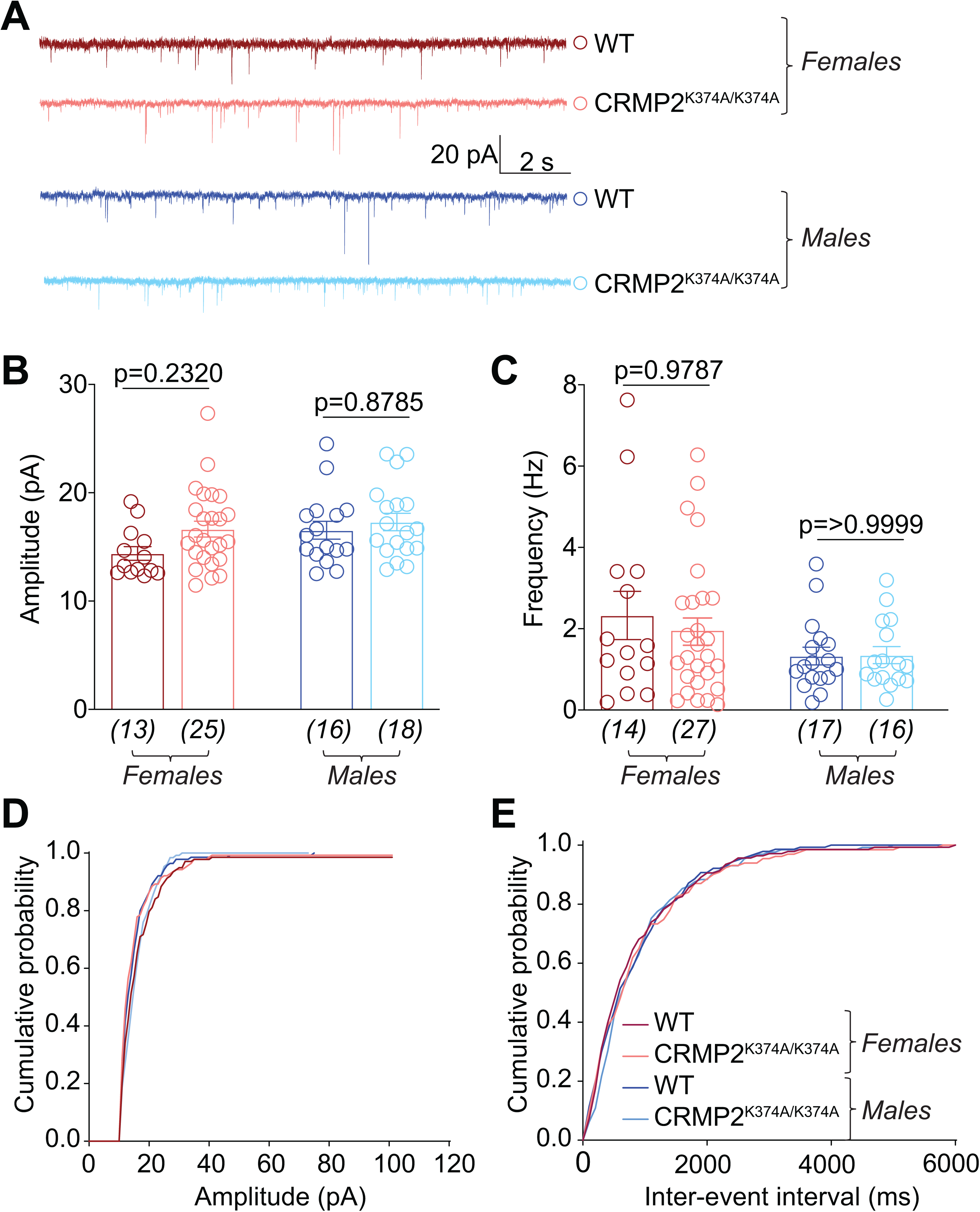
Excitatory neurotransmission is not affected in the lumbar dorsal horn of CRMP2^K374A/K374A^ knock-in mice. (**A**) Representative traces of cells from both sexes and genotypes. Bar graph with scatter plot showing the summary of amplitudes (**B**) and frequencies (**C**) of spontaneous excitatory post-synaptic currents (sEPSCs) for the indicated groups are shown. The cumulative probability of amplitude (**D**) and inter-event interval (**E**) are indicated. No significant change was observed in either parameter across any of the conditions tested. Data are shown as mean ± S.E.M., n=13-27 cells from at least 4 mice per experimental condition. p > 0.05, one-way ANOVA followed by Tukey’s post hoc test. Error bars indicate mean ± SEM. The experiments were conducted by investigators blinded to the genotype.

### 3.9 In vivo inhibition of voltage gated sodium channels in CRMP2^K374A/K374A^ mice

Our *in vitro* investigations showed that in females CRMP2^K374A/K374A^ mice, the interaction of NaV1.7 with CRMP2 was reduced, leading to decreased membrane localization and currents. To test if this regulation translates to a phenotype *in vivo*, we used the sodium channel opener veratridine [9], which causes all sodium channels to remain open upon repolarization following a step depolarization [8] and has been demonstrated to induce pain-like lifting and licking behaviors in mice [26]. Injecting veratridine (1 mg) in the paw of mice elicited a licking response that lasted for about 20-min. Consistent with our electrophysiology and imaging observations, in female CRMP2^K374A/K374A^ mice, we observed a significant decrease in paw licking time (**Figure 9A**). Male CRMP2^K374A/K374A^ mice showed no difference in paw licking time compared to wildtype mice (**Figure 9B**). These results indicated that the CRMP2^K374A/K374A^ mutation confers sex-specific regulation of NaV1.7 in vivo.

**Figure 9.**
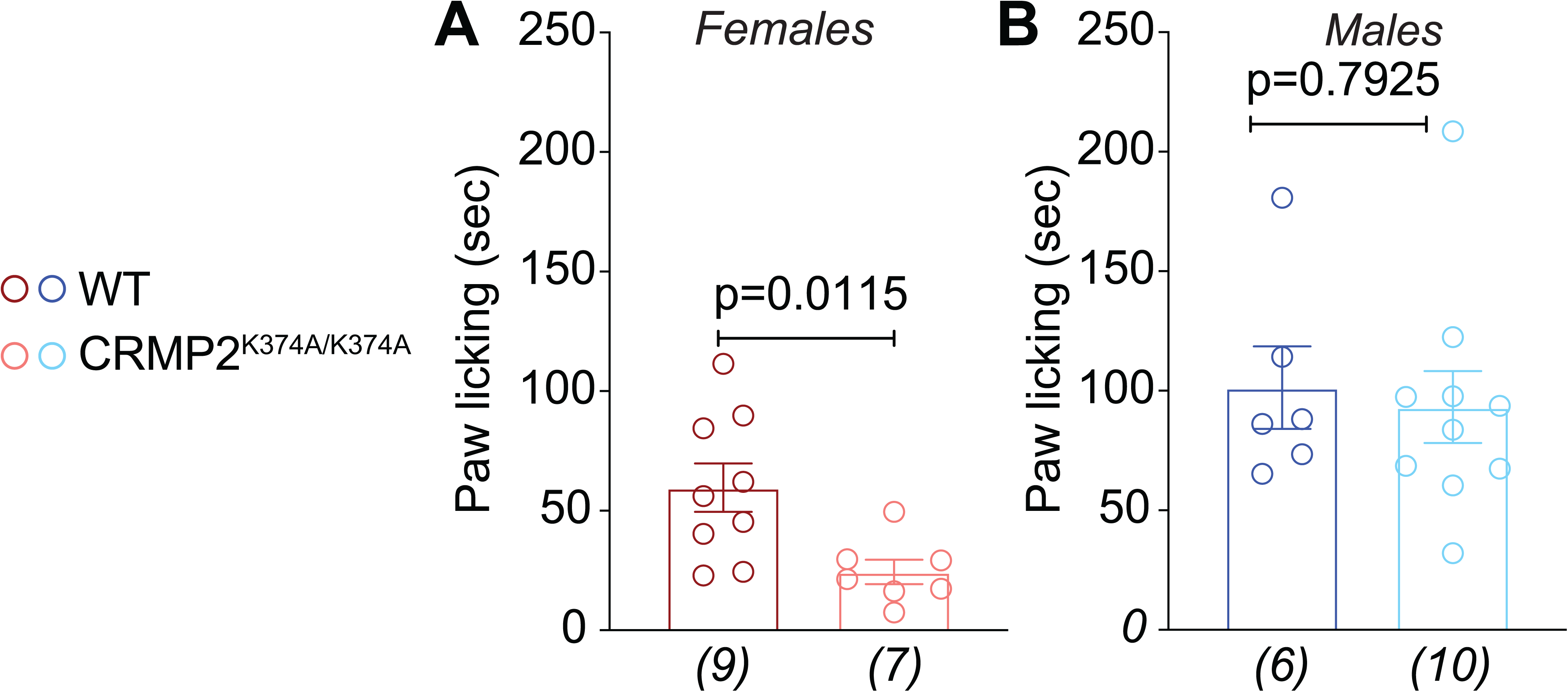
Female CRMP2^K374A/K374A^ knock-in mice have increased pain-like behaviors in response to injection of the sodium channel activator veratridine. Licking in the indicated groups of mice in response to an intraplantar dose of 1-mg veratridine. At this dose, veratridine caused a statistically significant increase in paw licking and lifting behaviors compared to saline. A change was seen in CRMP2^K374A/K374A^ female mice compared to wildtype counterparts (**A**), while no differences between genotypes were observed in males (**B**). See statistical analysis described in **Table 2**. Error bars indicate mean ± SEM. The experiments were conducted by investigators blinded to the genotype.

### 3.10 Effect of the CRMP2^K374A/K374A^ mutation on the development of chronic neuropathic pain

Thus far, a salient finding in our study is that female CRMP2^K374A/K374A^ mice have decreased NaV1.7 function *in vitro* and *in vivo*, while male CRMP2^K374A/K374A^ mice have NaV1.7 unaltered function. We previously found that in the spared nerve injury (SNI) model of chronic neuropathic pain, CRMP2 SUMOylation was upregulated in male rats [47]. Further, forced expression of a form of CRMP2 resistant to SUMOylation in the DRG and spinal cord of male rats reversed their SNI-induced mechanical allodynia [47]. Additionally, a ‘decoy’ peptide preventing CRMP2 SUMOylation reversed SNI-elicited allodynia in male rats [22]. Based on these observations, we posited that the CRMP2^K374A/K374A^ mice might show resistance to the development of mechanical allodynia after SNI. Therefore, performed SNI surgery in these mice and followed the development of mechanical allodynia for 35 days (**Figure 10**). In male and female WT mice, mechanical allodynia reached the lowest threshold at day 7 after surgery and lasted for the duration of the experiment (**Figure 10**). In female CRMP2^K374A/K374A^ mice, we found significantly higher paw withdrawal thresholds after SNI compared to WT mice (**Figure 10A, B**). While WT mice maintained a strong mechanical allodynia at 35 days after SNI, the paw withdrawal threshold in female CRMP2^K374A/K374A^ mice was restored back to its pre-surgery level (**Figure 10A**). Similarly, male CRMP2^K374A/K374A^ mice had mechanical allodynia after SNI that was less pronounced than in WT mice (**Figure 10C, D**). This allodynia almost completely resolved after 35 days (**Figure 10C, D**). Together, these results show for the first time that limiting CRMP2 SUMOylation can prevent the development of chronic allodynia after a neuropathic pain injury (SNI) in both male and female mice.

**Figure 10.**
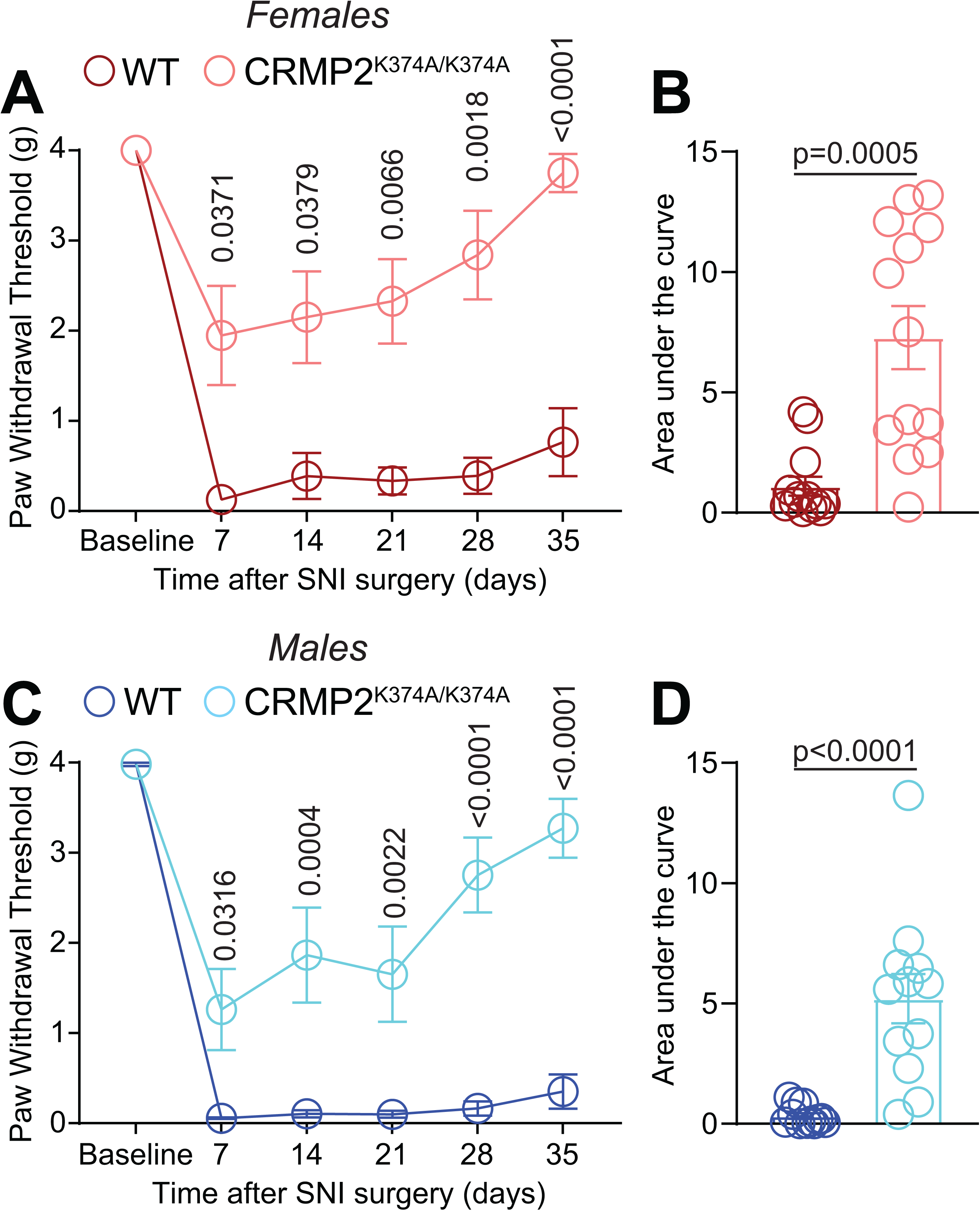
CRMP2^K374A/K374A^ knock-in mice do not develop persistent mechanical allodynia in the spared nerve injury (SNI) model of neuropathic pain. Paw withdrawal thresholds of age-matched and genotyped WT and CRMP2^K374A/K374A^ mice were measured at baseline and for five weeks following SNI. Post SNI, von Frey testing was confined to the sural nerve innervating region of the paw. Time course (**A** – *females*; **C** – *males*) and area under the curve (**B** – *females*; **D** – *males*) are shown. Area under the curve for paw withdrawal thresholds was derived using the trapezoid method. von Frey mechanical thresholds indicating that loss of CRMP2 SUMOylation prevented the development of mechanical allodynia after SNI in both male and female CRMP2^K374A/K374A^ mice. See statistical analysis described in **Table 2**. Error bars indicate mean ± SEM. The experiments were conducted by investigators blinded to the genotype.

### 3.11 Role of CRMP2 SUMOylation in non-chronic pain models

Because our findings support an unequivocal role for CRMP2 SUMOylation in chronic neuropathic pain (**Figure 10**), we next asked if the regulation of NaV1.7 by SUMOylated CRMP2 could be important for non-chronic pain. First, we used a model of inflammatory nociception with formalin as the chemical irritant. This model allows for the evaluation of two phases, a first phase related to direct activation of nociceptors followed by a second phase associated with spinal cord sensitization [58]. Following injection of a 2% formalin solution in the paw, all mice displayed licking behavior that lasted for 60 min. We quantified the paw licking time in bins of 5-minutes and found no effect of the CRMP2^K374A/K374A^ genotype in either sex on licking time (**Figure 11A-D**).

**Figure 11.**
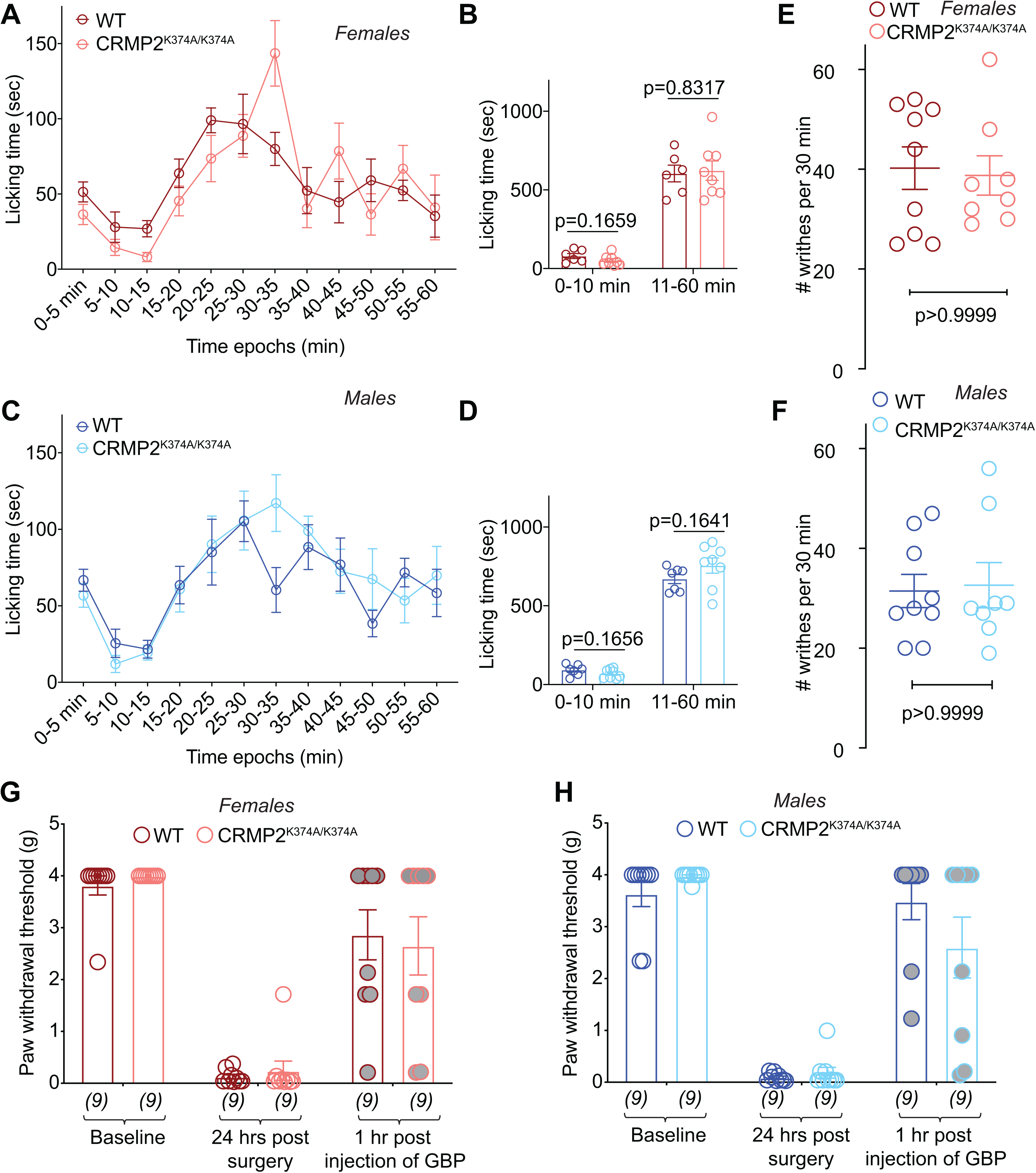
CRMP2^K374A/K374A^ knock-in mice do not display increased responsiveness to chemical, surgical, or visceral models of pain. Time course of number of flinches following subcutaneous (dorsal surface of the paw) injection of formalin (2.5% in 50 μl saline) in female (**A**) and male (**C**) wildtype (WT) and CRMP2^K374A/K374A^ mice. The total number of flinches in formalin-induced phase 1 (0-10 min) and phase 2 (11-60 min) (**B, D**). No significant differences were detected between the both sexes or genotypes. Writhing, characterized by abdominal stretching combined with an exaggerated extension of the hind limbs, was induced by intraperitoneal injection of acetic acid. No significant differences were detected between the both sexes or genotypes (**E, F**). Mice received a plantar incision on the left hind paw. Paw withdrawal thresholds were significantly decreased 24 hours after incision in both sexes and genotypes (**G, H**). Gabapentin (GBP), at a dose of 100 mg/kg (i.p.), reversed the mechanical allodynia 1-hour post administration. See statistical analysis described in **Table 2**. Error bars indicate mean ± SEM. The experiments were conducted by investigators blinded to the genotype.

Visceral pain is a modality reported to be independent of NaV1.7 [31], although adult-onset deletion of NaV1.7 using cre recombination conferred resistance to acetic acid induced pain [63]. We previously found that blocking CRMP2 SUMOylation with a SUMOylation motif decoy peptide had no effect on writhing induced by peritoneal acetic acid administration [22]. Here, we tested if CRMP2^K374A/K374A^ mice had altered sensitivity to visceral pain. No difference in the number of writhes elicited by an intraperitoneal injection of 1% acetic acid were found in CRMP2^K374A/K374A^ mice compared to wildtype mice of either sex (**Figure 11E, F**), showing that CRMP2 SUMOylation is not involved in visceral pain sensation.

Finally, we tested a model of post-surgical pain [59]. We previously reported that blunting CRMP2 phosphorylation or interfering with the CRMP2-CaV2.2 interaction could reverse mechanical allodynia following a paw incision [24; 46]. However, the role of CRMP2 SUMOylation in post-surgical pain was previously unknown. After subjecting mice to a paw incision and measuring allodynia at 24 hours post-operation, we did not find any significant effect on the CRMP2^K374A/K374A^ genotype (**Figure 11G, H**). As a positive control, we used gabapentin (100mg/kg, i.p.) [37], which reversed mechanical allodynia in these mice (**Figure 11G, H**). Altogether, our results highlight a role for CRMP2 SUMOylation in allodynia after neuropathic pain but not in non-chronic pain models.

## 4. Discussion

In this study, to elucidate the physiological role of SUMOylation of CRMP2 at Lys374, we newly generated CRMP2 knock-in mutant mice using CRISPR/Cas9 in which the Lys residue at 374 was replaced with Ala. For the first time, we demonstrate that SUMOylation of CRMP2 at Lys374 plays an essential role in the development of chronic neuropathic pain. Reduced thermal nociceptive processing and anxiety were other notable phenotypic traits exhibited by the knock-in mice. Furthermore, by analyzing sensory neurons from CRMP2^K374A/K374A^ mutant knock-in mice, we revealed dramatic regulation of NaV1.7 trafficking and activity in female mice; sex-specific regulation of NaV1.7 has never been previously reported. Without effects on other sensory neuron voltage-gated ion channels’ basal excitability, nor hippocampal and spinal cord plasticity (at both presynaptic and postsynaptic sites), these results establish CRMP2 SUMOylation as a selective, safe, and novel pharmacological target for pain.

A salient finding of this study is the female-specific regulation of NaV1.7 trafficking and currents in sensory neurons. While sexual dimorphism in pain behaviors has now been widely reported (e.g. [28; 40]), to our knowledge, no previous studies have reported evidence of sex-specific NaV1.7 regulation in neurons both *in vivo* and *in vitro*. A parsimonious explanation for this could lie in potential compromise of CRMP2’s ability to regulate NaV1.7 within males. Although previous studies have demonstrated a link between neurotrophins (e.g., BDNF [65] and NGF [45]) and CRMP2/NaV1.7 [25; 30; 33; 54; 66], adding these factors to DRGs from male CRMP2^K374A/K374A^ mice had no effect on NaV1.7. Within male mice DRGs, increased heterogeneity of NaV1.7 surface expression alongside more limited inhibition (53% in males vs. 76% in females) following NaV1.7-selective inhibitor PF05089771 application suggested that NaV1.7 trafficking and function may be less efficient compared to the parallel process in female CRMP2^K374A/K374A^ mice. Because NaV1.7 trafficking is clathrin dependent [17], we reasoned that blocking endocytosis (with Pitstop2) may enhance NaV1.7 currents in male CRMP2^K374A/K374A^ mice DRGs. While inhibiting clathrin assembly with Pitstop2 canceled the inhibition of sodium currents observed in female CRMP2^K374A/K374A^ mice DRGs, no effect was observed in either wildtype or CRMP2^K374A/K374A^ male mice DRGs; this excludes involvement of clathrin-mediated endocytosis as a possible explanation. Furthermore, the female-specific NaV1.7 regulation cannot be attributed to differences in CRMP2 itself, since CRMP2 expression and phosphorylation appear to be similar between sexes [60]. The extent of CRMP2 SUMOylation was unchanged but the binding between NaV1.7 and CRMP2 was lower in DRGs from male compared to female wildtype mice. These results point to CRMP2-independent trafficking for a pool of NaV1.7, which may be more prevalent in male mice. Another likely explanation for reduced NaV1.7-CRMP2 interaction amongst male mice DRGs, a yet unknown protein may compete with CRMP2 for the same binding site on NaV1.7. Although NaV1.7’s 267 member interactome in male mice was recently reported [10; 35], comparison with a similar dataset in female mice may be necessary to enable discovery of NaV1.7’s sex-specific trafficking mechanisms.

Alternatively, and not tested here, CRMP2 SUMOylation may differentially control the *recycling* of NaV1.7 to account for the sex-specific difference in NaV1.7 trafficking. Our previous study demonstrated increased colocalization of the recycling endosome protein Rab11 with NaV1.7 when CRMP2 was deSUMOylated in DRGs from female rat [17]. Inhibition of channel recycling was recently reported as a mechanism for down regulation of a calcium channel [67]. CRMP2 can control vesicular trafficking via its interaction with the Molecule Interacting with CasL Like (MICAL1) protein [62] – a protein controlling slow endocytic recycling of endosomes [27].

Accumulating work from our laboratory has established CRMP2 SUMOylation as an important contributor to electrogenesis and neuropathic pain [17; 47]. We have also solved the crystal structure of CRMP2 [18; 23] and identified key residues in the SUMOylation interface that are amenable to targeting by small molecules. The generation of the CRMP2^K374A/K374A^ mouse obviates reliance on overexpression of mutant proteins (e.g., SUMO-null CRMP2) which, if produced at high levels, may be toxic or induce the unfolded protein response [7; 61], which could affect NaV1.7 trafficking indirectly. Of the CRMP2 transgenic mice reported thus far (i.e., global [55] or brain-specific [74] CRMP2 knockout; cyclin dependent kinase 5-phosphorylation-null CRMP2^S522A/S522A^ mutant knock-in mice [71]; O-GlcNAcylation-null CRMP2^S517A/S517A^ mutant knock-in mice [53]), none have investigated females. Mice with a global deletion of CRMP2 had deficits in contextual and social learning and decreased anxiety [55]. Additionally, the brain-specific CRMP2 knockout mice exhibited behavioral deficits in locomotor activity, sensorimotor gating, social behavior and spatial learning and memory as well as hippocampal synaptic dysfunction [74]. By contrast, CRMP2^K374A/K374A^ mice displayed unaffected theta-burst stimulation-induced long-term potentiation. Both the input–output curve and paired-pulse ratio showed no difference, indicating unperturbed presynaptic function. These results imply that contextual learning is likely to be unaffected in the CRMP2^K374A/K374A^ mice. In the marble-burying and nestlet shredding tests, which evaluate repetitive, compulsive-like behaviors related to autism-like OCD [3; 57], no abnormal phenotype was noted in CRMP2^K374A/K374A^ mice. A decreased anxiety phenotype was seen in both female and male CRMP2^K374A/K374A^ mice in our study, which is consistent with results from male global CRMP2 knockout mice [55] and CRMP2^S522A/S522A^ mutant knock-in mice [71]. The general lack of behavioral impairments observed in our CRMP2^K374A/K374A^ mice indicates that CRMP2 SUMOylation may not be critical for learning and memory and autism-like OCD behaviors.

An exciting discovery in this study is that the establishment of chronic neuropathic pain is dependent on CRMP2 SUMOylation. Both female and male CRMP2^K374A/K374A^ mice did not develop allodynia as markedly as wildtype mice. This is in accordance with our previous observations showing that targeting CRMP2 SUMOylation, either genetically [47] or with a peptide [22], reverses nerve-injury induced mechanical allodynia in rats. Another remarkable observation was that, in both female and male CRMP2^K374A/K374A^ mice, nerve-injury induced mechanical allodynia was completely resolved within 5 weeks of the injury. Therefore, we infer that preventing CRMP2 SUMOylation has a disease-modifying effect. Given previously reported upregulation of CRMP2 SUMOylation in male rats alongside these findings [47], CRMP2 SUMOylation warrants consideration as an objective biomarker to potentially guide drug development and clinical practice for pain management. CRMP2^K374A/K374A^ mice had unaltered behavioral responses compared to their wildtype counterparts in inflammatory, visceral and post-surgical pain models. While these results may at first appear contradictory, as other strategies targeting CRMP2 have demonstrated efficacy [5; 46], they instead indicate a selective involvement of CRMP2 SUMOylation in the pathophysiology of chronic neuropathic pain.

The generation of CRMP2^K374A/K374A^ mice has been predicated on the consistent demonstration by our laboratory [17; 22; 24; 47] and others [14; 21; 72] of a key role of CRMP2 in regulation of nociceptive ion channels and pain. In particular, we reported that elimination of the CRMP2 SUMO modification site in heterologous cells was sufficient to selectively decrease trafficking and activity of NaV1.7, but not NaV1.1 or NaV1.3 or NaV1.5 channels [19]. In rat and human DRGs, tetrodotoxin (TTX)-resistant (NaV1.8, NaV1.9) as well as TTX-sensitive (NaV1.2, and NaV1.6) currents were also unaffected upon loss of CRMP2 SUMOylation [17; 19]. Using the CRMP2^K374A/K374A^ mice allowed us to characterize the biological consequences of the selective regulation of NaV1.7 by CRMP2 SUMOylation. Prior efforts to target NaV1.7 for pain relief focused on development of direct blockers of the channel but have remained unsuccessful [20; 36; 43; 64]. Some reasons (among those disclosed) for why these NaV1.7-targeting drugs failed include issues with central nervous system penetration [1; 2; 64], lack of selectivity (e.g. of Biogen’s Vixotrigine [1]), inadequacy of pain models [64], and inadequate degree of block leading to a lack of action potential inhibition [20]. Global or nociceptor-specific deletion of NaV1.7 abolished thermal nociception as well as pain behaviors, notably in inflammatory (formalin) and post-surgical pain models [26; 56; 63]. In CRMP2^K374A/K374A^ mice, we observed: (*i*) a significant increase in paw withdrawal latency to noxious thermal stimulation (52°C) in both female and male CRMP2^K374A/K374A^ mice, which was more pronounced in female CRMP2^K374A/K374A^ mice at 48°C, (*ii*) no change across genotypes or sexes in the tail-flick assay which is a reflexive nociceptive test independent of higher brain functions, and (*iii*) responsivity in inflammatory post-surgical pain models. Nevertheless, female CRMP2^K374A/K374A^ mice exhibit reduced pain responses to the NaV1.7-activator veratridine, thus showing that the channel is indeed inhibited by this mechanism *in vivo*. Our experiments with the NaV1.7-selective compound PF05089771 demonstrate that ∼50% of NaV1.7 is inhibited in DRGs from female CRMP2^K374A/K374A^ mice. This incomplete block of NaV1.7 may contribute to the lack of phenotype in thermal, inflammatory, and post-surgical pain in female CRMP2^K374A/K374A^ mice. Altogether, these findings are congruent with the hypothesis that therapeutic targeting may need to achieve complete block of NaV1.7 to achieve analgesia.

In summary, the actions of CRMP2 SUMOylation appear to be peripherally restricted and important for the pathophysiology of chronic neuropathic pain. To this end, the CRMP2^K374A/K374A^ mouse offers a unique platform for studying CRMP2 SUMOylation in membrane trafficking and cell biology as well as assessing the specificity of future drug candidates targeting CRMP2 SUMOylation to modulate NaV1.7.

## ACKNOWLEDGEMENTS

The pX330-U6-Chimeric_BB-CBh-hSpCas9 was a gift from Professor Feng Zhang (Addgene plasmid # 42230). This work was supported by National Institutes of Health (NIH) awards (NS098772 from the National Institute of Neurological Disorders and Stroke and DA042852 from the National Institute on Drug Abuse to R.K). L.A.C. is supported by the Duke University School of Medicine Medical Scientist Training Program T32GM007171. The CRMP2 transgenic mice were created by the Genetically Engineered Mouse Model core at the University of Arizona by Drs. Thomas C. Deutschman and Teodora G Georgieva via an Institutional Mouse Precision Modeling grant to R.K.. Partial support for animal husbandry reported in this publication was provided by the Experimental Mouse Shared Resource (EMSR) core which was funded by the National Cancer Institute of the NIH under award number P30 CA023074.

