## Supplementary Table for "Studies on CRMP2 SUMOylation-deficient transgenic mice identify sex-specific NaV1.7 regulation in the pathogenesis of chronic neuropathic pain"

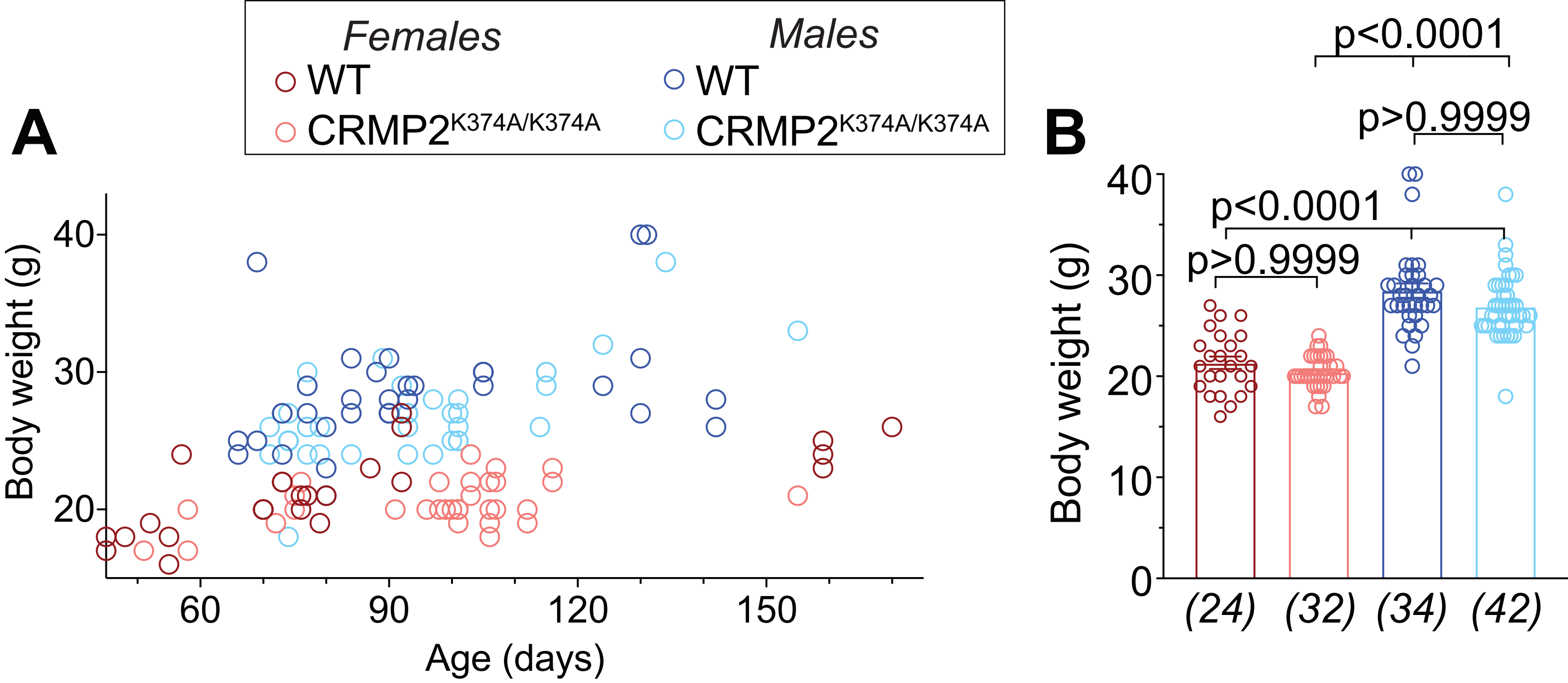
Females
WT
CRMP2K374A/K374A
Males
WT
CRMP2K374A/K374A
A
B
Body weight (g)
Age (days)
40
30
20
60
90
120
150
10
0
p<0.0001
p>0.9999
p<0.0001
p>0.9999
(24)
(32)
(34)
(42)

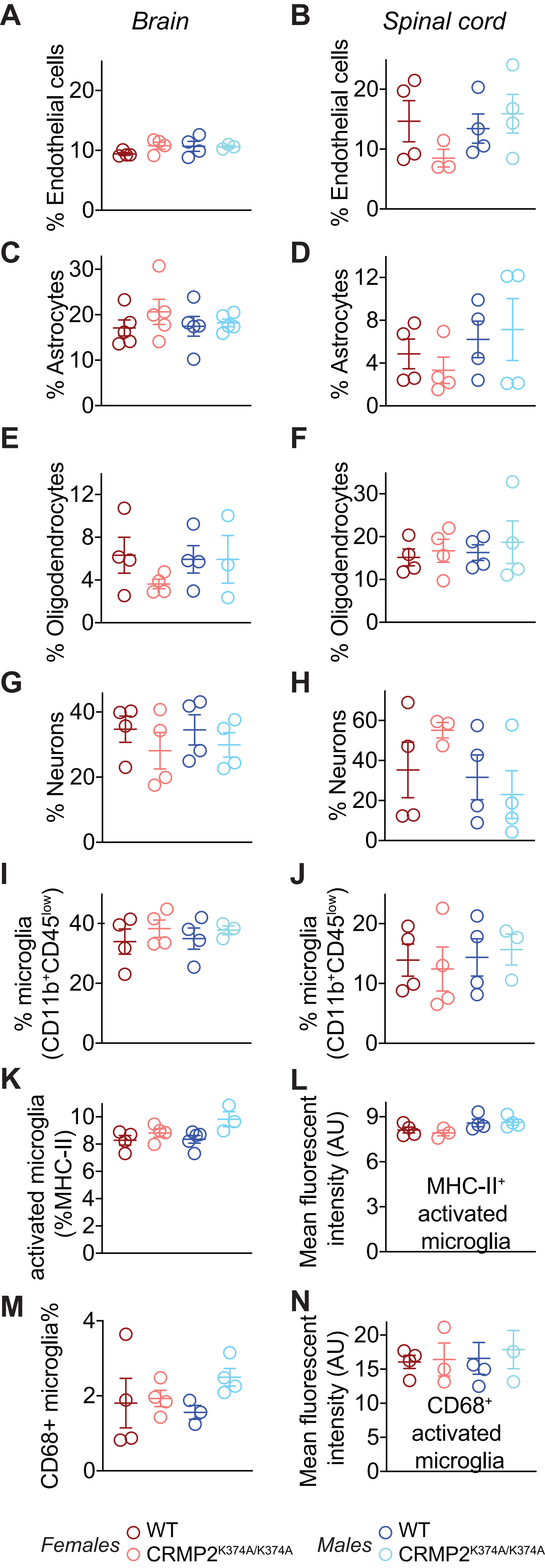

A
Brain
% Endothelial cells
B
Spinal cord
% Endothelial cells
C
% Astrocytes
D
% Astrocytes
E
% Oligodendrocytes
F
% Oligodendrocytes
G
% Neurons
H
% Neurons
I
% microglia (CD11b+CD45low)
J
% microglia (CD11b+CD45low)
K
activated microglia (%MHC-II)
L
Mean fluorescent intensity (AU)
MHC-II+ activated microglia
M
CD68+ microglia%
N
Mean fluorescent intensity (AU)
CD68+ activated microglia
Females
WT
CRMP2K374A/K374A
Males
WT
CRMP2K374A/K374A

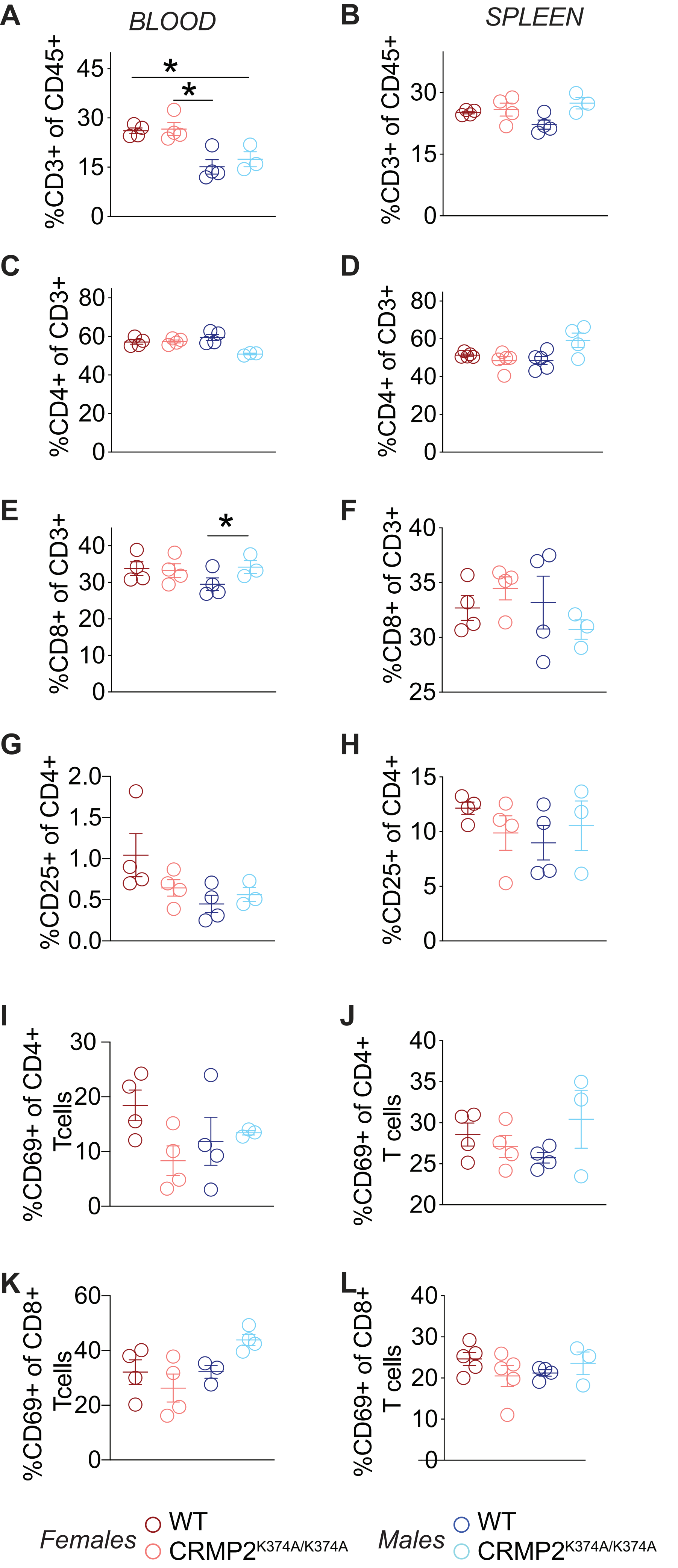
A
BLOOD
B
SPLEEN
%CD3+ of CD45+
C
D
%CD4+ of CD3+
E
F
%CD8+ of CD3+
G
H
%CD25+ of CD4+
I
%CD69+ of CD4+ Tcells
J
%CD69+ of CD4+ T cells
K
%CD69+ of CD8+ Tcells
L
%CD69+ of CD8+ T cells
Females
WT
CRMP2K374A/K374A
Males
WT
CRMP2K374A/K374A

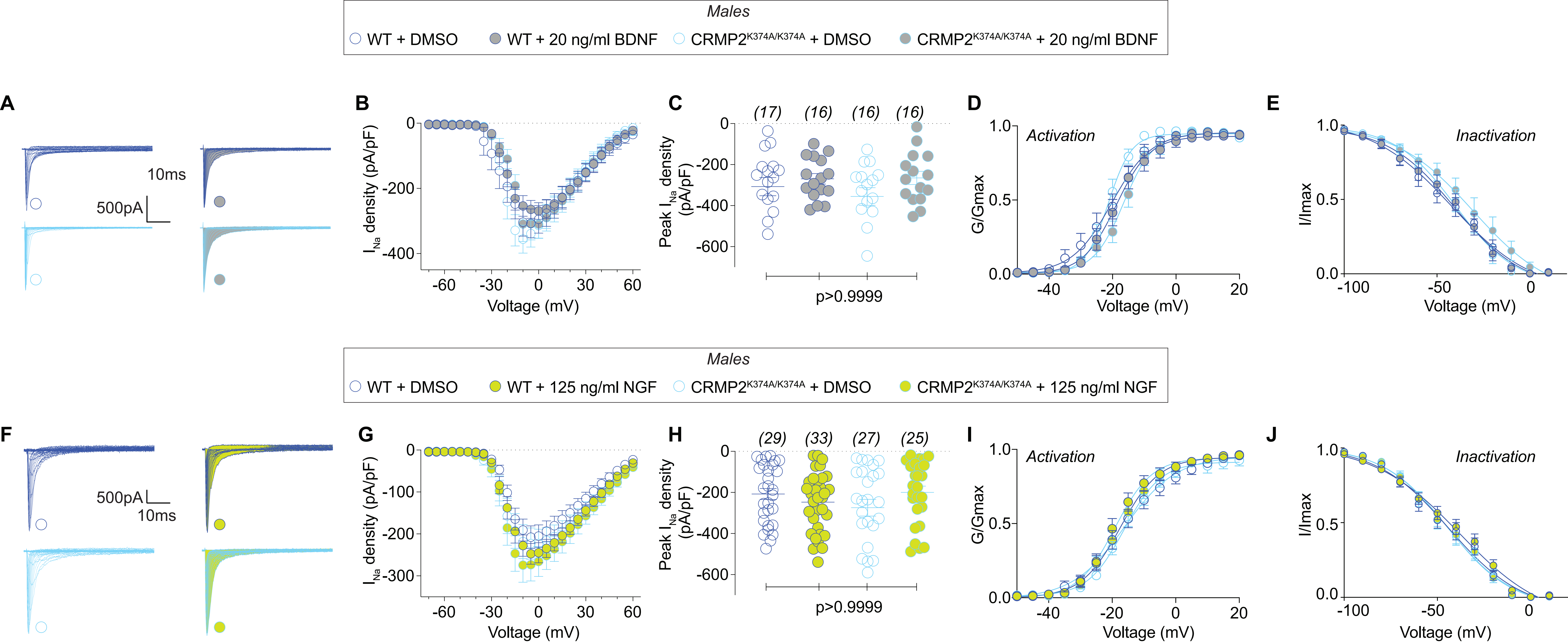
Males
WT + DMSO
WT + 20 ng/ml BDNF
CRMP2K374A/K374A + DMSO
CRMP2K374A/K374A + 20 ng/ml BDNF
A
10ms
500pA
B
INa density (pA/pF)
Voltage (mV)
C
(17) (16) (16) (16)
Peak INa density (pA/pF)
p>0.9999
D
Activation
G/Gmax
Voltage (mV)
E
Inactivation
I/Imax
Voltage (mV)
Males
WT + DMSO
WT + 125 ng/ml NGF
CRMP2K374A/K374A + DMSO
CRMP2K374A/K374A + 125 ng/ml NGF
F
500pA
10ms
G
INa density (pA/pF)
Voltage (mV)
H
(29) (33) (27) (25)
Peak INa density (pA/pF)
p>0.9999
I
Activation
G/Gmax
Voltage (mV)
J
Inactivation
I/Imax
Voltage (mV)

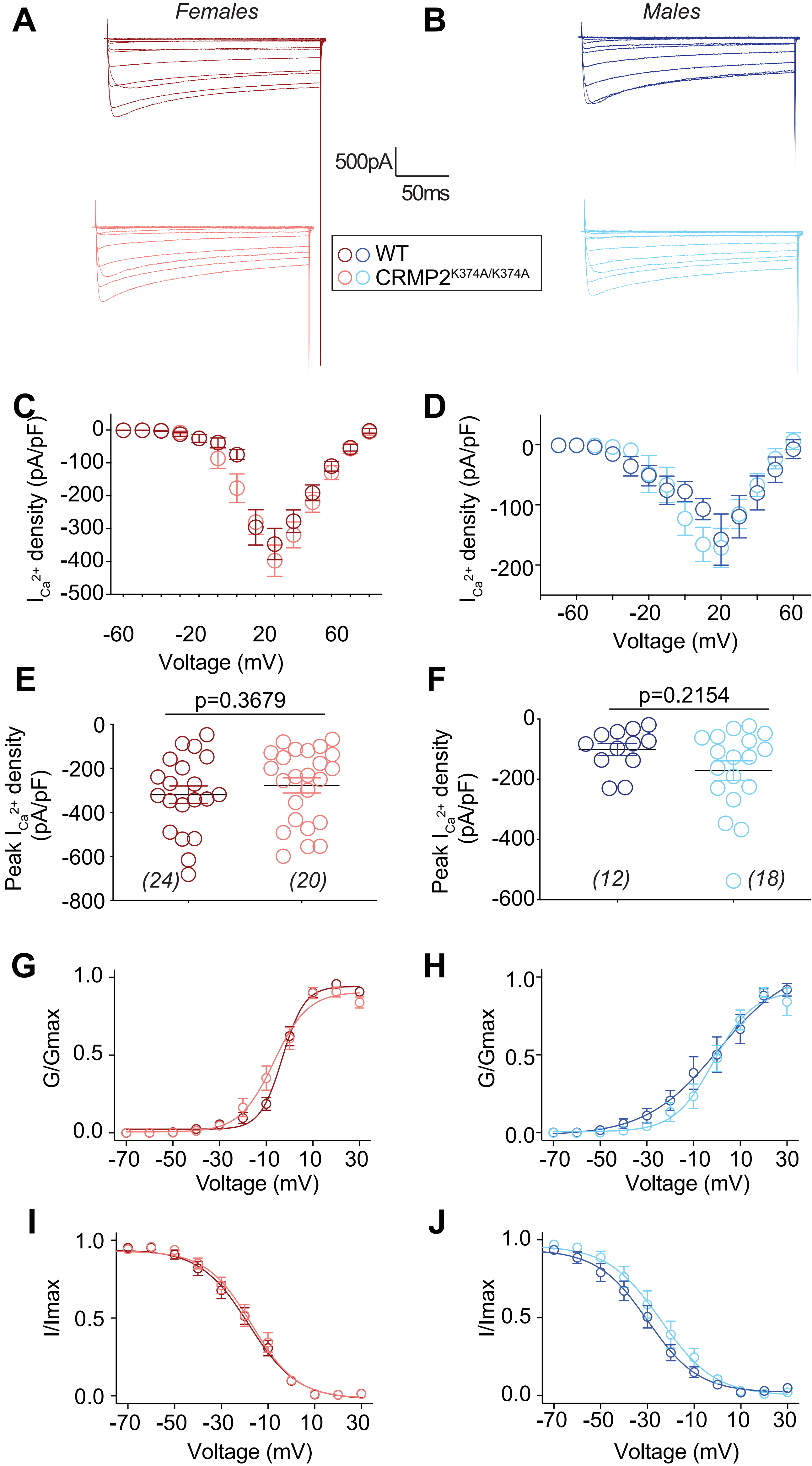

A
Females
B
Males
500pA
50ms
WT
CRMP2K374A/K374A
C
D
I$_{Ca}$$^{2+}$ density (pA/pF)
Voltage (mV)
E
F
p=0.3679
p=0.2154
Peak I$_{Ca}$$^{2+}$ density (pA/pF)
(24)
(20)
(12)
(18)
G
H
G/Gmax
I
J
I/Imax

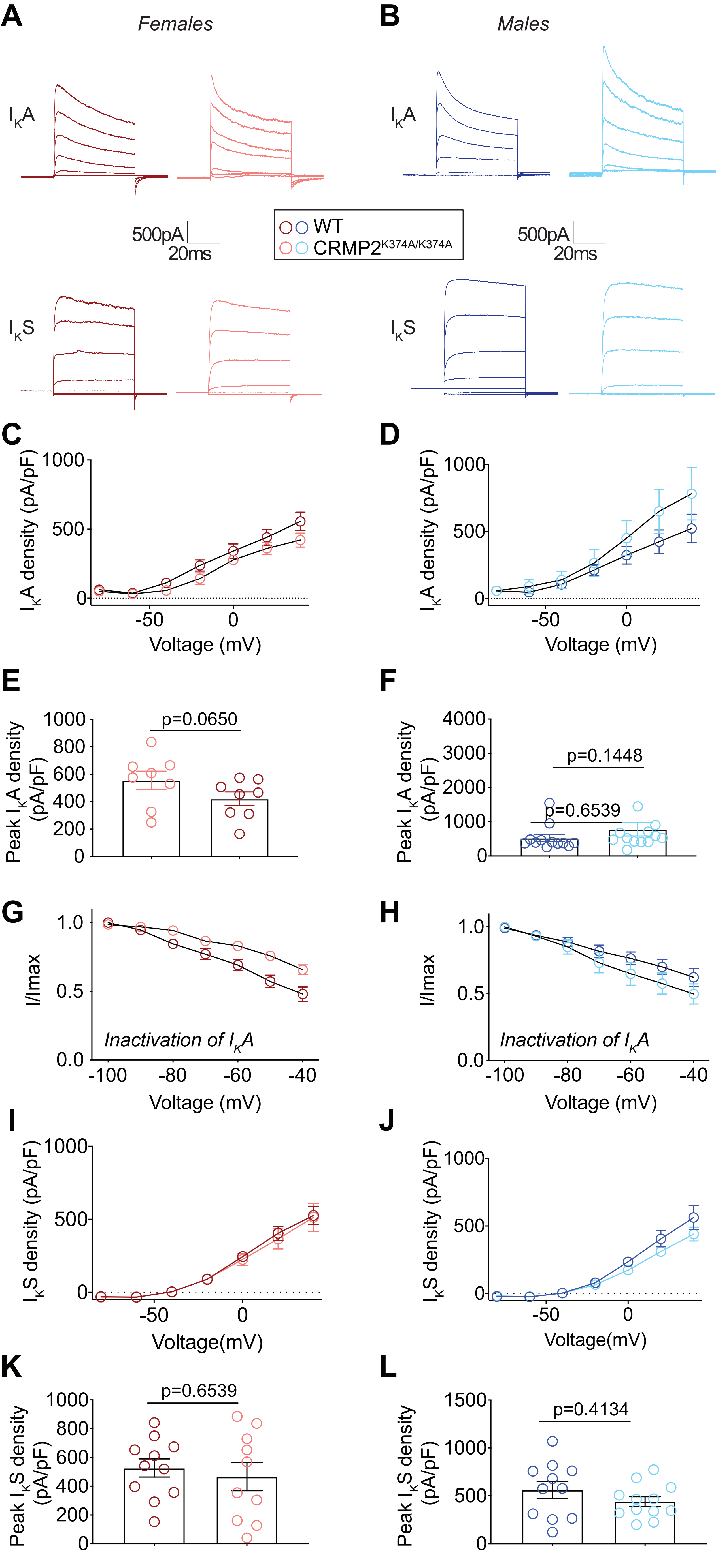
A
Females
B
Males
$I_KA$
$I_KA$
500pA
20ms
WT
CRMP2$^{K374A/K374A}$
500pA
20ms
$I_KS$
$I_KS$
C
$I_KA$ density (pA/pF)
1000
500
0
-50
0
Voltage (mV)
D
$I_KA$ density (pA/pF)
1000
500
0
-50
0
Voltage (mV)
E
Peak $I_KA$ density (pA/pF)
1000
800
600
400
200
0
p=0.0650
F
Peak $I_KA$ density (pA/pF)
4000
3000
2000
1000
0
p=0.1448
p=0.6539
G
I/Imax
1.0
0.5
0.0
Inactivation of $I_KA$
-100
-80
-60
-40
Voltage (mV)
H
I/Imax
1.0
0.5
0.0
Inactivation of $I_KA$
-100
-80
-60
-40
Voltage (mV)
I
$I_KS$ density (pA/pF)
1000
500
0
-50
0
Voltage(mV)
J
$I_KS$ density (pA/pF)
1000
500
0
-50
0
Voltage(mV)
K
Peak $I_KS$ density (pA/pF)
1000
800
600
400
200
0
p=0.6539
L
Peak $I_KS$ density (pA/pF)
1500
1000
500
0
p=0.4134

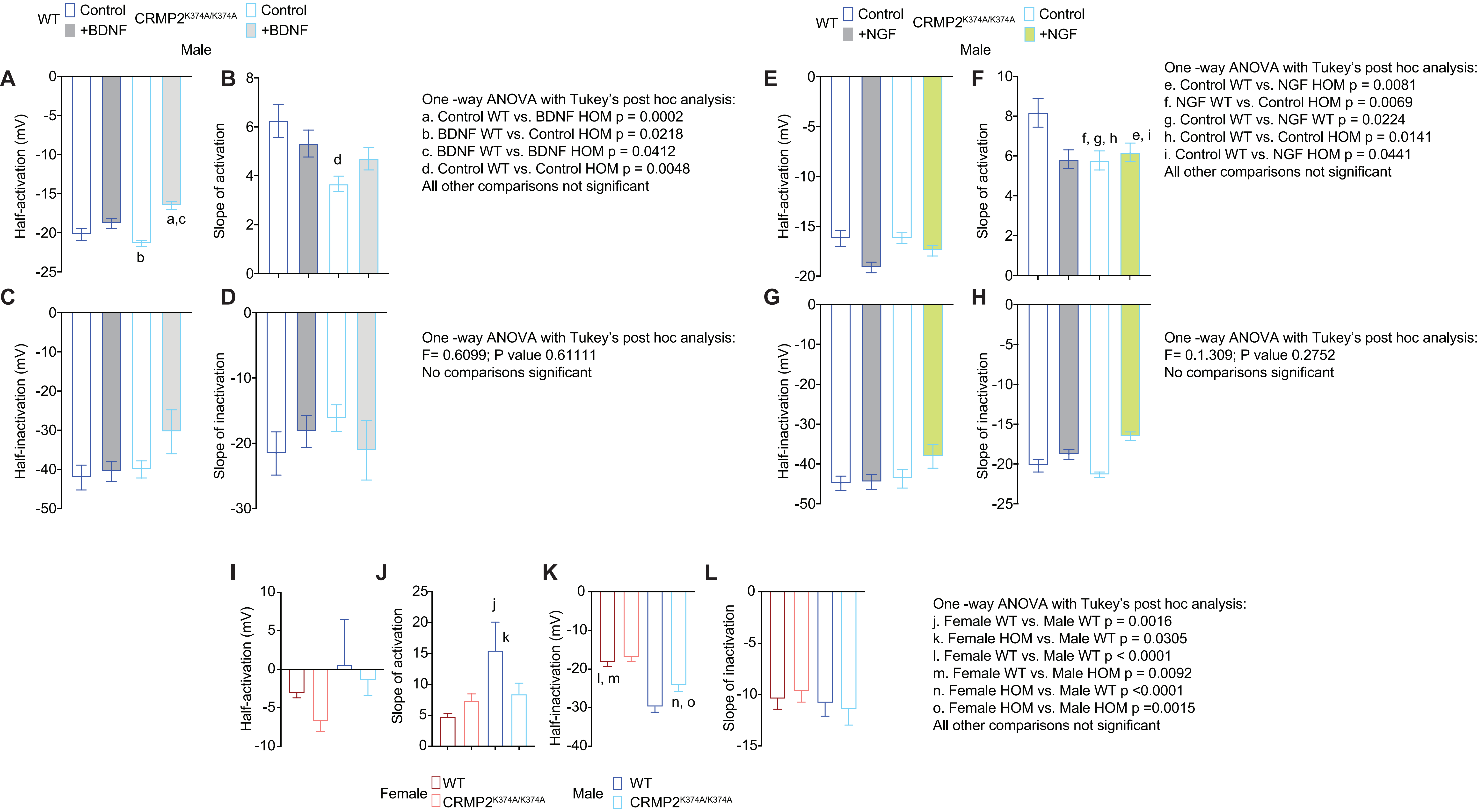
WT Control +BDNF CRMP2K374A/K374A Control +BDNF
Male
A
Half-activation (mV)
a,c
b
B
Slope of activation
d
One -way ANOVA with Tukey's post hoc analysis:
a. Control WT vs. BDNF HOM p = 0.0002
b. BDNF WT vs. Control HOM p = 0.0218
c. BDNF WT vs. BDNF HOM p = 0.0412
d. Control WT vs. Control HOM p = 0.0048
All other comparisons not significant
C
Half-inactivation (mV)
D
Slope of inactivation
One -way ANOVA with Tukey's post hoc analysis:
F= 0.6099; P value 0.61111
No comparisons significant
WT Control +NGF CRMP2K374A/K374A Control +NGF
Male
E
Half-activation (mV)
F
Slope of activation
f, g, h
e, i
One -way ANOVA with Tukey's post hoc analysis:
e. Control WT vs. NGF HOM p = 0.0081
f. NGF WT vs. Control HOM p = 0.0069
g. Control WT vs. NGF WT p = 0.0224
h. Control WT vs. Control HOM p = 0.0141
i. Control WT vs. NGF HOM p = 0.0441
All other comparisons not significant
G
Half-inactivation (mV)
H
Slope of inactivation
One -way ANOVA with Tukey's post hoc analysis:
F= 0.1.309; P value 0.2752
No comparisons significant
I
Half-activation (mV)
J
Slope of activation
j
k
K
Half-inactivation (mV)
l, m
n, o
L
Slope of inactivation
One -way ANOVA with Tukey's post hoc analysis:
j. Female WT vs. Male WT p = 0.0016
k. Female HOM vs. Male WT p = 0.0305
l. Female WT vs. Male WT p < 0.0001
m. Female WT vs. Male HOM p = 0.0092
n. Female HOM vs. Male WT p <0.0001
o. Female HOM vs. Male HOM p =0.0015
All other comparisons not significant
Female WT CRMP2K374A/K374A
Male WT CRMP2K374A/K374A

### Supplementary Figure Legends

**Supplementary Figure 1. Characterization of CRMP2$^{K374A/K374A}$ knock-in mice reveals no changes in bodyweight phenotype.** (**A**) Comparison of the body weights of WT and CRMP2$^{K374A/K374A}$ knock-in mice between 45–170 days old. (**B**) Comparison of the body weights of all WT and CRMP2$^{K374A/K374A}$ knock-in mice. Number of mice weighed are as indicated. There were no significant differences in body weight within sexes of CRMP2$^{K374A/K374A}$ mice compared to their WT littermates (45–170 days old or 6–22 weeks of age) – Females: 21.0 ± 0.61 g in WT [n = 24]; 20.0 ± 0.29 g in CRMP2$^{K374A/K374A}$ [n = 32]; Males: 28.0 ± 0.71 g in WT [n = 34]; 27.0 ± 0.49 g in CRMP2$^{K374A/K374A}$ [n = 42] (p>0.9999, ANOVA with a Kruskal-Wallis post hoc test). Male mice, irrespective of genotype, were larger than female mice (p<0.0001, ANOVA with a Kruskal-Wallis post hoc test).

**Supplementary Figure 2. Cellular makeup of the brain and spinal cord are not affected by CRMP2 mutation or sex.** Female and male wildtype (WT) and CRMP2$^{K374A/K374A}$ knock-in mice were used for this experiment. At necropsy the brains (*left column of scatter plots*) and spinal cords (*right column of scatter plots*) were collected, dissociated, stained and analyzed by flow cytometry. (**A**, **B**) Endothelial cells are characterized as CD31$^{+}$. (**C**, **D**) Astrocytes were characterized as GLAST$^{+}$. (**E**, **F**) Oligodendrocytes were characterized as O4$^{+}$. (**G**, **H**) Neurons were characterized as CD45$^{-}$CD11b$^{-}$CD31$^{-}$GLAST$^{-}$O4$^{-}$. The percentage of microglia, characterized were as CDllb$^{+}$CD45$^{low}$, was measured in the brain (**I**). Activation in brain microglia was measured as %MHC-II+ (**K**) and %CD68+ (**M**). Microglia in the spinal cord (**J**) were also characterized as CDllb$^{+}$CD45$^{low}$. However, activation in this population was measured by mean fluorescence intensity (MFI) of the MHC-II (**L**) and CD68 (**N**) signal. There were no significant differences in cell types within and across sexes or genotypes. See statistical analysis described in **Supplementary Table 1**.

**Supplementary Figure 3. T cell composition is affected by both sex and CRMP2 mutation.** White blood cells and splenic cells were isolated from whole blood and spleens at necropsy and stained for T cell markers (CD3, CD4, CD8). Samples were then analyzed by flow cytometry. The percentage of T cells in the blood (**A**) and spleen (**B**) were calculated. The T cell composition of CD4$^{+}$ T cells in the blood (**C**) and spleen (**D**) were measured as a percentage of CD3$^{+}$ cells. A similar analysis was done for CD8+ T cells in the blood (**E**) and spleen (**F**). The % of CD25+ CD4+ T cells was measured as a percentage of CD4+ T cells in the blood (**G**) and spleen (**H**). % CD4+ and CD8+ T cell activation (CD69+) was measured in the blood (**I, K**) and spleen (**J, L**). See statistical analysis described in **Supplementary Table 1**.

**Supplementary Figure 4. Nerve growth factor (NGF) or brain derived neurotrophic factor (BDNF) supplementation does not affect NaV1.7 currents in DRGs from male WT and CRMP2$^{K374A/K374A}$ knock-in mice.** In some experiments, currents were recorded after an overnight application with the indicated concentrations of NGF or BDNF. Representative traces of Na$^{+}$ currents from DRG sensory neurons isolated from male WT and CRMP2$^{K374A/K374A}$ knock-in mice (A – *BDNF*, F – *NGF*). Summary of current-voltage curves (**B** – *BDNF*, **G** – *NGF*) and normalized peak (**C** – *BDNF*, **H** – *NGF*) currents (picoAmperes/picoFarads, pA/pF) from small-to-medium diameter DRG neurons of WT and CRMP2$^{K374A/K374A}$ knock-in mice (n = 16-33).. There were no differences in sodium currents between genotypes and sexes in the presence or absence of the factors. Boltzmann fits for normalized conductance $G/G_{max}$ voltage relations for voltage dependent activation (**D** – *BDNF*, **I** – *NGF*) and inactivation (**E** – *BDNF*, **J** – *NGF*) of the sensory neurons were unchanged across any condition. Half-maximal activation and inactivation ($V_{1/2}$) and slope values (*k*) for activation and inactivation are presented in **Supplementary Figure 7**. Error bars indicate mean ± SEM.

**Supplementary Figure 5. Calcium currents are not affected by loss of CRMP2 SUMOylation.** Representative traces of total $Ca^{2+}$ currents from DRG sensory neurons isolated from female (**A**) and male (**B**) WT and CRMP2$^{K374A/K374A}$ knock-in mice. Currents were evoked by 200ms pulse between –70 and +60 mV. Summary of the normalized (pA/pF) total calcium current density versus voltage relationship (**C** – *female*, **D** – *male*) and peak (**E** – *female*, **F** – *male*) total $Ca^{2+}$ current density at +10 mV (mean ± SEM) from DRG sensory neurons treated as indicated. There were no differences in calcium currents between genotypes and sexes (n=12-24 per condition). Boltzmann fits for normalized conductance $G/G_{max}$ voltage relations for voltage dependent activation (**G** – *female*, **H** – *male*) and inactivation (**I** – *female*, **J** – *male*) of sensory neurons treated were unchanged across any condition. Half-maximal activation and inactivation ($V_{1/2}$) and slope values ($k$) for activation and inactivation are presented in **Supplementary Figure 7**. Error bars indicate mean ± SEM.

**Supplementary Figure 6. Potassium currents are not affected by loss of CRMP2 SUMOylation.** Representative family of $I_K$A and $I_K$S currents recorded from DRG neurons isolated from female (**A**) and male (**B**) WT and CRMP2$^{K374A/K374A}$ knock-in mice. Current density–voltage relationship of $I_K$A from DRGs isolated from female (**C**) and male (**D**) WT and CRMP2$^{K374A/K374A}$ knock-in mice. Peak current density of $I_K$A from female (**E**) and male (**F**) mice from each genotype. Inactivation of potassium channels from DRGs isolated from female (**G**) and male (**H**) WT and CRMP2$^{K374A/K374A}$ knock-in mice. There were no differences in potassium currents or biophysical properties between genotypes and sexes. n=8-13 cells/condition. Current density–voltage relationship of $I_K$S from DRGs isolated from female (**I**) and male (**J**) WT and CRMP2$^{K374A/K374A}$ knock-in mice. Peak current density of $I_K$S from female (**K**) and male (**L**) mice from each genotype (n=10-11 cells/condition). P values are as indicated (Unpaired Student's test with Mann-Whitney post-hoc). Error bars indicate mean ± SEM.

**Supplementary Figure 7. Half-activation and inactivation and slope values for sodium and calcium currents.** Half-maximal activation and inactivation ($V_{1/2}$) and slope values ($k$) were calculated from the Boltzmann fits for normalized conductance $G/G_{max}$ voltage relations for voltage dependent activation and inactivation.

**Supplementary Table 1. Statistical analyses of immune cell profiling.**

| Figure panel | Assay | Statistical test; findings | Post-hoc analysis (adjusted p-values) | Number of subjects |
|---|---|---|---|---|
| Supplementary Figure 2A | %Endothelial cells in brain | Kruskal Wallis test P = 0.3329 | Dunn's multiple comparisons test<br>WT females vs. CRMP2$^{K374A/K374A}$females: p =0.4919<br>WT females vs. WT males: p >0.9999<br>WT females vs. CRMP2$^{K374A/K374A}$males: p =0.9423<br>CRMP2$^{K374A/K374A}$females vs. WT males: p >0.9999<br>CRMP2$^{K374A/K374A}$females vs. CRMP2$^{K374A/K374A}$males: p>0.9999<br>WT females vs. CRMP2$^{K374A/K374A}$females: p >0.9999<br>WT males vs. CRMP2$^{K374A/K374A}$males: p >0.9999 | WT females: n = 4<br>CRMP2$^{K374A/K374A}$females: n = 4<br>WT males: n = 4<br>CRMP2$^{K374A/K374A}$males: n = 3 |
| Supplementary Figure 2B | %Endothelial cells in spinal cord | Kruskal Wallis test P = 0.3048 | Dunn's multiple comparisons test<br>WT females vs. CRMP2$^{K374A/K374A}$females: p =0.9423<br>WT females vs. WT males: p >0.9999<br>WT females vs. CRMP2$^{K374A/K374A}$males: p >0.9999<br>CRMP2$^{K374A/K374A}$females vs. WT males: p =0.8201<br>CRMP2$^{K374A/K374A}$females vs. CRMP2$^{K374A/K374A}$males: p=0.3823<br>WT females vs. CRMP2$^{K374A/K374A}$females: p >0.9999<br>WT males vs. CRMP2$^{K374A/K374A}$males: p >0.9999 | WT females: n = 4<br>CRMP2$^{K374A/K374A}$females: n = 3<br>WT males: n = 4<br>CRMP2$^{K374A/K374A}$males: n = 4 |
| Supplementary Figure 2C | %Astrocytes in brain | Kruskal Wallis test P = 0.6581 | Dunn's multiple comparisons test<br>WT females vs. CRMP2$^{K374A/K374A}$females: p >0.9999<br>WT females vs. WT males: p >0.9999<br>WT females vs. CRMP2$^{K374A/K374A}$males: p >0.9999 | WT females: n = 5<br>CRMP2$^{K374A/K374A}$females: n = 5<br>WT males: n =5<br>CRMP2$^{K374A/K374A}$males: n = 5 |

| | | | | |
|---|---|---|---|---|
| | | | $CRMP2^{K374A/K374A}$females vs. WT males: p >0.9999<br>$CRMP2^{K374A/K374A}$females vs. $CRMP2^{K374A/K374A}$males: p>0.9999<br>WT females vs. $CRMP2^{K374A/K374A}$females: p >0.9999<br>WT males vs. $CRMP2^{K374A/K374A}$males: p >0.9999 | |
| Supplementary Figure 2D | %Astrocytes in spinal cord | Kruskal Wallis test P = 0.6559 | Dunn's multiple comparisons test<br>WT females vs. $CRMP2^{K374A/K374A}$females: p >0.9999<br>WT females vs. WT males: p >0.9999<br>WT females vs. $CRMP2^{K374A/K374A}$males: p >0.9999<br>$CRMP2^{K374A/K374A}$females vs. WT males: p >0.9999<br>$CRMP2^{K374A/K374A}$females vs. $CRMP2^{K374A/K374A}$males: p>0.9999<br>WT females vs. $CRMP2^{K374A/K374A}$females: p >0.9999<br>WT males vs. $CRMP2^{K374A/K374A}$males: p >0.9999 | WT females: n = 4<br>$CRMP2^{K374A/K374A}$females: n = 4<br>WT males: n = 4<br>$CRMP2^{K374A/K374A}$males: n =4 |
| Supplementary Figure 2E | %oligodendrocytes in brain | Kruskal Wallis test P = 0.4339 | Dunn's multiple comparisons test<br>WT females vs. $CRMP2^{K374A/K374A}$females: p =0.6831<br>WT females vs. WT males: p >0.9999<br>WT females vs. $CRMP2^{K374A/K374A}$males: p >0.9999<br>$CRMP2^{K374A/K374A}$females vs. WT males: p >0.9999<br>$CRMP2^{K374A/K374A}$females vs. $CRMP2^{K374A/K374A}$males: p>0.9999<br>WT females vs. $CRMP2^{K374A/K374A}$females: p >0.9999<br>WT males vs. $CRMP2^{K374A/K374A}$males: p >0.9999 | WT females: n = 4<br>$CRMP2^{K374A/K374A}$females: n = 4<br>WT males: n = 4<br>$CRMP2^{K374A/K374A}$males: n = 3 |
| Supplementary Figure 2F | %oligodendrocytes in spinal cord | Kruskal Wallis test P = 0.9712 | Dunn's multiple comparisons test | WT females: n = 4<br>$CRMP2^{K374A/K374A}$females: n = 4 |

| | | | | |
|---|---|---|---|---|
| | | | WT females vs. CRMP2$^{K374A/K374A}$females: p >0.9999<br>WT females vs. WT males: p >0.9999<br>WT females vs. CRMP2$^{K374A/K374A}$males: p >0.9999<br>CRMP2$^{K374A/K374A}$females vs. WT males: p >0.9999<br>CRMP2$^{K374A/K374A}$females vs. CRMP2$^{K374A/K374A}$males: p>0.9999<br>WT females vs. CRMP2$^{K374A/K374A}$females: p >0.9999<br>WT males vs. CRMP2$^{K374A/K374A}$males: p >0.9999 | WT males: n = 4<br>CRMP2$^{K374A/K374A}$males: n = 4 |
| Supplementary Figure 2G | %neurons in brain | Kruskal Wallis test P = 0.4784 | Dunn's multiple comparisons test<br>WT females vs. CRMP2$^{K374A/K374A}$females: p >0.9999<br>WT females vs. WT males: p >0.9999<br>WT females vs. CRMP2$^{K374A/K374A}$males: p >0.9999<br>CRMP2$^{K374A/K374A}$females vs. WT males: p =0.9495<br>CRMP2$^{K374A/K374A}$females vs. CRMP2$^{K374A/K374A}$males: p>0.9999<br>WT females vs. CRMP2$^{K374A/K374A}$females: p >0.9999<br>WT males vs. CRMP2$^{K374A/K374A}$males: p >0.9999 | WT females: n = 4<br>CRMP2$^{K374A/K374A}$females: n = 4<br>WT males: n = 4<br>CRMP2$^{K374A/K374A}$males: n = 4 |
| Supplementary Figure 2H | %neurons in spinal cord | Kruskal Wallis test P = 0.2583 | Dunn's multiple comparisons test<br>WT females vs. CRMP2$^{K374A/K374A}$females: p >0.9999<br>WT females vs. WT males: p =0.3236<br>WT females vs. CRMP2$^{K374A/K374A}$males: p >0.9999<br>CRMP2$^{K374A/K374A}$females vs. WT males: p =0.6128<br>CRMP2$^{K374A/K374A}$females vs. CRMP2$^{K374A/K374A}$males: p>0.9999 | WT females: n = 4<br>CRMP2$^{K374A/K374A}$females: n = 3<br>WT males: n = 4<br>CRMP2$^{K374A/K374A}$males: n = 4 |

| | | | | |
|---|---|---|---|---|
| | | | WT females vs. CRMP2$^{K374A/K374A}$females: p >0.9999<br>WT males vs. CRMP2$^{K374A/K374A}$males: p >0.9999 | |
| Supplementary Figure 2I | %microglia in brain | Kruskal Wallis test P = 0.8347 | Dunn's multiple comparisons test<br>WT females vs. CRMP2$^{K374A/K374A}$females: p >0.9999<br>WT females vs. WT males: p >0.9999<br>WT females vs. CRMP2$^{K374A/K374A}$males: p >0.9999<br>CRMP2$^{K374A/K374A}$females vs. WT males: p >0.9999<br>CRMP2$^{K374A/K374A}$females vs. CRMP2$^{K374A/K374A}$males: p>0.9999<br>WT females vs. CRMP2$^{K374A/K374A}$females: p >0.9999<br>WT males vs. CRMP2$^{K374A/K374A}$males: p >0.9999 | WT females: n = 4<br>CRMP2$^{K374A/K374A}$females: n = 4<br>WT males: n = 4<br>CRMP2$^{K374A/K374A}$males: n = 3 |
| Supplementary Figure 2J | %microglia in spinal cord | Kruskal Wallis test P = 0.8433 | Dunn's multiple comparisons test<br>WT females vs. CRMP2$^{K374A/K374A}$females: p >0.9999<br>WT females vs. WT males: p >0.9999<br>WT females vs. CRMP2$^{K374A/K374A}$males: p >0.9999<br>CRMP2$^{K374A/K374A}$females vs. WT males: p >0.9999<br>CRMP2$^{K374A/K374A}$females vs. CRMP2$^{K374A/K374A}$males: p>0.9999<br>WT females vs. CRMP2$^{K374A/K374A}$females: p >0.9999<br>WT males vs. CRMP2$^{K374A/K374A}$males: p >0.9999 | WT females: n = 4<br>CRMP2$^{K374A/K374A}$females: n = 4<br>WT males: n = 4<br>CRMP2$^{K374A/K374A}$males: n = 3 |
| Supplementary Figure 2K | %MHC-II+ microglia in brain | Kruskal Wallis test P = 0.0458 | Dunn's multiple comparisons test<br>WT females vs. CRMP2$^{K374A/K374A}$females: p >0.9999<br>WT females vs. WT males: p >0.9999 | WT females: n = 4<br>CRMP2$^{K374A/K374A}$females: n = 4<br>WT males: n = 5<br>CRMP2$^{K374A/K374A}$males: n = 3 |

| | | | | |
|---|---|---|---|---|
| | | | WT females vs. CRMP2$^{K374A/K374A}$males: p =0.1292<br>CRMP2$^{K374A/K374A}$females vs. WT males: p >0.9999<br>CRMP2$^{K374A/K374A}$females vs. CRMP2$^{K374A/K374A}$males: p>0.9999<br>WT females vs. CRMP2$^{K374A/K374A}$females: p >0.9999<br>WT males vs. CRMP2$^{K374A/K374A}$males: p =0.0973 | |
| Supplementary Figure 2L | MHC-II (MFI) of microglia in spinal cord | Kruskal Wallis test P = 0.1008 | Dunn's multiple comparisons test<br>WT females vs. CRMP2$^{K374A/K374A}$females: p >0.9999<br>WT females vs. WT males: p >0.9999<br>WT females vs. CRMP2$^{K374A/K374A}$males: p =0.7985<br>CRMP2$^{K374A/K374A}$females vs. WT males: p =0.3823<br>CRMP2$^{K374A/K374A}$females vs. CRMP2$^{K374A/K374A}$males: p=0.1908<br>WT females vs. CRMP2$^{K374A/K374A}$females: p >0.9999<br>WT males vs. CRMP2$^{K374A/K374A}$males: p >0.9999 | WT females: n = 4<br>CRMP2$^{K374A/K374A}$females: n = 3<br>WT males: n = 4<br>CRMP2$^{K374A/K374A}$males: n = 4 |
| Supplementary Figure 2M | %CD68+ microglia in brain | Kruskal Wallis test P = 0.1906 | Dunn's multiple comparisons test<br>WT females vs. CRMP2$^{K374A/K374A}$females: p >0.9999<br>WT females vs. WT males: p >0.9999<br>WT females vs. CRMP2$^{K374A/K374A}$males: p =0.5334<br>CRMP2$^{K374A/K374A}$females vs. WT males: p >0.9999<br>CRMP2$^{K374A/K374A}$females vs. CRMP2$^{K374A/K374A}$males: p>0.9999<br>WT females vs. CRMP2$^{K374A/K374A}$females: p >0.9999<br>WT males vs. CRMP2$^{K374A/K374A}$males: p =0.2637 | WT females: n = 4<br>CRMP2$^{K374A/K374A}$females: n = 4<br>WT males: n = 3<br>CRMP2$^{K374A/K374A}$males: n = 4 |

| | | | | |
|---|---|---|---|---|
| Supplementary Figure 2N | CD68 (MFI) of microglia in spinal cord | Kruskal Wallis test P = 0.6976 | Dunn's multiple comparisons test<br>WT females vs. CRMP2$^{K374A/K374A}$females: p >0.9999<br>WT females vs. WT males: p >0.9999<br>WT females vs. CRMP2$^{K374A/K374A}$males: p >0.9999<br>CRMP2$^{K374A/K374A}$females vs. WT males: p >0.9999<br>CRMP2$^{K374A/K374A}$females vs. CRMP2$^{K374A/K374A}$males: p>0.9999<br>WT females vs. CRMP2$^{K374A/K374A}$females: p >0.9999<br>WT males vs. CRMP2$^{K374A/K374A}$males: p >0.9999 | WT females: n = 4<br>CRMP2$^{K374A/K374A}$females: n = 4<br>WT males: n = 4<br>CRMP2$^{K374A/K374A}$males: n = 3 |

| | | | | |
|---|---|---|---|---|
| Supplementary Figure 3A | %CD3 of CD45+ in Blood | Kruskal Wallis test P = 0.0009 | Dunn's multiple comparisons test<br>WT females vs. CRMP2$^{K374A/K374A}$females: p >0.9999<br>WT females vs. WT males: p =0.0431<br>WT females vs. CRMP2$^{K374A/K374A}$males: p = 0.4261<br>CRMP2$^{K374A/K374A}$females vs. WT males: p = 0.0431<br>CRMP2$^{K374A/K374A}$females vs. CRMP2$^{K374A/K374A}$males: p =0.4261<br>WT females vs. CRMP2$^{K374A/K374A}$females: p >0.9999<br>WT males vs. CRMP2$^{K374A/K374A}$males: p >0.9999 | WT females: n = 4<br>CRMP2$^{K374A/K374A}$females: n = 4<br>WT males: n = 4<br>CRMP2$^{K374A/K374A}$males: n = 3 |
| Supplementary Figure 3B | %CD3 of CD45+ in Spleen | Kruskal Wallis test P = 0.1337 | Dunn's multiple comparisons test<br>WT females vs. CRMP2$^{K374A/K374A}$females: p >0.9999<br>WT females vs. WT males: p =0.0431<br>WT females vs. CRMP2$^{K374A/K374A}$males: p = 0.4261<br>CRMP2$^{K374A/K374A}$females vs. WT males: p = 0.5813<br>CRMP2$^{K374A/K374A}$females vs. CRMP2$^{K374A/K374A}$males: p >0.9999 | WT females: n = 4<br>CRMP2$^{K374A/K374A}$females: n = 4<br>WT males: n = 4<br>CRMP2$^{K374A/K374A}$males: n = 3 |

| | | | | |
|---|---|---|---|---|
| | | | WT females vs. CRMP2$^{K374A/K374A}$females: p >0.9999<br>WT males vs. CRMP2$^{K374A/K374A}$males: p=0.1488 | |
| Supplementary Figure 3C | %CD4 of CD3+ in Blood | Kruskal Wallis test P = 0.0304 | Dunn's multiple comparisons test<br>WT females vs. CRMP2$^{K374A/K374A}$females: p >0.9999<br>WT females vs. WT males: p >0.9999<br>WT females vs. CRMP2$^{K374A/K374A}$males: p = 0.4739<br>CRMP2$^{K374A/K374A}$females vs. WT males: p >0.9999<br>CRMP2$^{K374A/K374A}$females vs. CRMP2$^{K374A/K374A}$males: p=0.2027<br>WT females vs. CRMP2$^{K374A/K374A}$females: p >0.9999<br>WT males vs. CRMP2$^{K374A/K374A}$males: p=0.0406 | WT females: n = 4<br>CRMP2$^{K374A/K374A}$females: n = 4<br>WT males: n = 4<br>CRMP2$^{K374A/K374A}$males: n = 3 |
| Supplementary Figure 3D | %CD4 of CD3+ in Spleen | Kruskal Wallis test P = 0.0894 | Dunn's multiple comparisons test<br>WT females vs. CRMP2$^{K374A/K374A}$females: p >0.9999<br>WT females vs. WT males: p >0.9999<br>WT females vs. CRMP2$^{K374A/K374A}$males: p >0.9999<br>CRMP2$^{K374A/K374A}$females vs. WT males: p >0.9999<br>CRMP2$^{K374A/K374A}$females vs. CRMP2$^{K374A/K374A}$males: p=0.1851<br>WT females vs. CRMP2$^{K374A/K374A}$females: p =0.6937<br>WT males vs. CRMP2$^{K374A/K374A}$males: p=0.3092 | WT females: n = 5<br>CRMP2$^{K374A/K374A}$females: n = 5<br>WT males: n = 5<br>CRMP2$^{K374A/K374A}$males: n = 4 |
| Supplementary Figure 3E | %CD8 of CD3+ in Blood | Kruskal Wallis test P = 0.3621 | Dunn's multiple comparisons test<br>WT females vs. CRMP2$^{K374A/K374A}$females: p >0.9999<br>WT females vs. WT males: p =0.7985 | WT females: n = 4<br>CRMP2$^{K374A/K374A}$females: n = 4<br>WT males: n = 4<br>CRMP2$^{K374A/K374A}$males: n = 3 |

| | | | | |
|---|---|---|---|---|
| | | | WT females vs. $CRMP2^{K374A/K374A}$males: p >0.9999<br>$CRMP2^{K374A/K374A}$females vs. WT males: p =0.9284<br>$CRMP2^{K374A/K374A}$females vs. $CRMP2^{K374A/K374A}$males: p>0.9999<br>WT females vs. $CRMP2^{K374A/K374A}$females: p >0.9999<br>WT males vs. $CRMP2^{K374A/K374A}$males: p=0.7822 | |
| Supplementary Figure 3F | %CD8 of CD3+ in Spleen | Kruskal Wallis test P = 0.5315 | Dunn's multiple comparisons test<br>WT females vs. $CRMP2^{K374A/K374A}$females: p >0.9999<br>WT females vs. WT males: p >0.9999<br>WT females vs. $CRMP2^{K374A/K374A}$males: p >0.9999<br>$CRMP2^{K374A/K374A}$females vs. WT males: p >0.9999<br>$CRMP2^{K374A/K374A}$females vs. $CRMP2^{K374A/K374A}$males: p=0.7457<br>WT females vs. $CRMP2^{K374A/K374A}$females: p >0.9999<br>WT males vs. $CRMP2^{K374A/K374A}$males: p >0.9999 | WT females: n = 4<br>$CRMP2^{K374A/K374A}$females: n = 4<br>WT males: n = 4<br>$CRMP2^{K374A/K374A}$males: n = 3 |
| Supplementary Figure 3G | %CD25 of CD4+ in Blood | Kruskal Wallis test P = 0.0901 | Dunn's multiple comparisons test<br>WT females vs. $CRMP2^{K374A/K374A}$females: p =0.9262<br>WT females vs. WT males: p =0.948<br>WT females vs. $CRMP2^{K374A/K374A}$males: p =0.5663<br>$CRMP2^{K374A/K374A}$females vs. WT males: p >0.9999<br>$CRMP2^{K374A/K374A}$females vs. $CRMP2^{K374A/K374A}$males: p>0.9999<br>WT females vs. $CRMP2^{K374A/K374A}$females: p >0.9999<br>WT males vs. $CRMP2^{K374A/K374A}$males: p >0.9999 | WT females: n = 4<br>$CRMP2^{K374A/K374A}$females: n = 4<br>WT males: n = 4<br>$CRMP2^{K374A/K374A}$males: n = 3 |

| | | | | |
|---|---|---|---|---|
| Supplementary Figure 3H | %CD25 of CD4+ in Spleen | Kruskal Wallis test<br>P = 0.5618 | Dunn's multiple comparisons test<br>WT females vs. CRMP2$^{K374A/K374A}$females: p >0.9999<br>WT females vs. WT males: p >0.9999<br>WT females vs. CRMP2$^{K374A/K374A}$males: p >0.9999<br>CRMP2$^{K374A/K374A}$females vs. WT males: p >0.9999<br>CRMP2$^{K374A/K374A}$females vs. CRMP2$^{K374A/K374A}$males: p>0.9999<br>WT females vs. CRMP2$^{K374A/K374A}$females: p >0.9999<br>WT males vs. CRMP2$^{K374A/K374A}$males: p >0.9999 | WT females: n = 4<br>CRMP2$^{K374A/K374A}$females: n = 4<br>WT males: n = 4<br>CRMP2$^{K374A/K374A}$males: n = 3 |
| Supplementary Figure 3I | %CD69 of CD4+ in Blood | Kruskal Wallis test<br>P = 0.1663 | Dunn's multiple comparisons test<br>WT females vs. CRMP2$^{K374A/K374A}$females: p =0.2390<br>WT females vs. WT males: p =0.4919<br>WT females vs. CRMP2$^{K374A/K374A}$males: p >0.9999<br>CRMP2$^{K374A/K374A}$females vs. WT males: p >0.9999<br>CRMP2$^{K374A/K374A}$females vs. CRMP2$^{K374A/K374A}$males: p>0.9999<br>WT females vs. CRMP2$^{K374A/K374A}$females: p >0.9999<br>WT males vs. CRMP2$^{K374A/K374A}$males: p >0.9999 | WT females: n = 4<br>CRMP2$^{K374A/K374A}$females: n = 4<br>WT males: n = 4<br>CRMP2$^{K374A/K374A}$males: n = 3 |
| Supplementary Figure 3J | %CD69 of CD4+ in Spleen | Kruskal Wallis test<br>P = 0.5724 | Dunn's multiple comparisons test<br>WT females vs. CRMP2$^{K374A/K374A}$females: p >0.9999<br>WT females vs. WT males: p >0.9999<br>WT females vs. CRMP2$^{K374A/K374A}$males: p >0.9999<br>CRMP2$^{K374A/K374A}$females vs. WT males: p >0.9999<br>CRMP2$^{K374A/K374A}$females vs. CRMP2$^{K374A/K374A}$males: p>0.9999 | WT females: n = 4<br>CRMP2$^{K374A/K374A}$females: n = 4<br>WT males: n = 4<br>CRMP2$^{K374A/K374A}$males: n = 3 |

| | | | | |
|---|---|---|---|---|
| | | | WT females vs. CRMP2$^{K374A/K374A}$females: p >0.9999<br>WT males vs. CRMP2$^{K374A/K374A}$males: p >0.9999 | |
| Supplementary Figure 3K | %CD69 of CD8+ in Blood | Kruskal Wallis test P = 0.0185 | Dunn's multiple comparisons test<br>WT females vs. CRMP2$^{K374A/K374A}$females: p =0.0339<br>WT females vs. WT males: p =0.2572<br>WT females vs. CRMP2$^{K374A/K374A}$males: p =0.4141<br>CRMP2$^{K374A/K374A}$females vs. WT males: p >0.9999<br>CRMP2$^{K374A/K374A}$females vs. CRMP2$^{K374A/K374A}$males: p>0.9999<br>WT females vs. CRMP2$^{K374A/K374A}$females: p >0.9999<br>WT males vs. CRMP2$^{K374A/K374A}$males: p >0.9999 | WT females: n = 4<br>CRMP2$^{K374A/K374A}$females: n = 4<br>WT males: n = 3<br>CRMP2$^{K374A/K374A}$males: n = 4 |
| Supplementary Figure 3L | %CD69 of CD8+ in Spleen | Kruskal Wallis test P = 0.3578 | Dunn's multiple comparisons test<br>WT females vs. CRMP2$^{K374A/K374A}$females: p >0.9999<br>WT females vs. WT males: p =0.5708<br>WT females vs. CRMP2$^{K374A/K374A}$males: p >0.9999<br>CRMP2$^{K374A/K374A}$females vs. WT males: p >0.9999<br>CRMP2$^{K374A/K374A}$females vs. CRMP2$^{K374A/K374A}$males: p>0.9999<br>WT females vs. CRMP2$^{K374A/K374A}$females: p >0.9999<br>WT males vs. CRMP2$^{K374A/K374A}$males: p >0.9999 | WT females: n = 5<br>CRMP2$^{K374A/K374A}$females: n = 4<br>WT males: n = 4<br>CRMP2$^{K374A/K374A}$males: n =3 |
